# *Citrobacter rodentium* infection reveals preserved intestinal resilience in aged intestine

**DOI:** 10.64898/2026.07.27.740925

**Authors:** Priyanka Biswas, Dharitri Chaudhuri, Vishwas Mishra, Zuza Kozik, Julia Sanchez-Garrido, Harini Ramini, Jyoti S. Chaudhary, Sandhya S. Visweswariah, Gad Frankel

**Affiliations:** Department of Life Sciences, Imperial, London SW7 2AZ, UK; Department of Developmental Biology and Genetics, Indian Institute of Science, Bengaluru, India; Functional Proteomics Group, Chester Beatty Laboratories, Institute of Cancer Research, London, SW3 6JB, United Kingdom

**Keywords:** Ageing, *Citrobacter rodentium*, type III secretion system effectors, infectious colitis, Intestinal resilience

## Abstract

Ageing is generally associated with increased susceptibility to enteric infection and is widely considered to reflect progressive deterioration of intestinal function. However, whether increased susceptibility represents complete loss of intestinal resilience or a distinct host response to infection remains unclear. Using *Citrobacter rodentium* (CR) infection as a model of infectious colitis, we investigated how ageing shapes epithelial responses to enteric bacterial infection. Infection of aged mice with wild type CR (CR_WT_) resulted greater bacterial colonisation, deeper crypt localisation, impaired fluid-ion homeostasis and more severe colitis than young mice. Conversely, infection of aged mice with CR_M12_, a CR strain lacking 12 type III secretion system effectors, resulted in reduced epithelial colonisation, preservation of epithelial barrier function and fluid-ion homeostasis, restrained inflammatory responses, and markedly lower disease severity than CR_WT_ infection. Deep quantitative proteomic analysis demonstrated that these divergent outcomes were accompanied by distinct adaptive molecular programmes within the aged intestinal epithelium. Together, our findings demonstrate that although aged mice exhibit increased susceptibility to enteric infection, the aged intestine retains the ability to mount distinct epithelial, inflammatory and molecular responses to pathogens with different virulence repertoires. Thus, increased susceptibility in aged mice should not be interpreted as complete loss of intestinal resilience.

## Introduction

Ageing represents a distinct physiological state characterised by widespread dysregulation of innate and adaptive immune responses, altered gut microbiota composition, changes in the intestinal stem cell function, and compromised epithelial barrier integrity^1–5^. These changes collectively reshape intestinal immune and epithelial homeostasis, increasing susceptibility to infection^6^. Consistently, aged mice exhibit heightened disease severity following infections with *Salmonella enterica* serovar Typhimurium^7,8^, influenza virus^9^, and *Candida albicans*^10^, typically linked to impaired immune function and cytokine responses^11^. Moreover, age-related changes in epithelial composition, mitochondrial function, and tight junction integrity further contribute to barrier vulnerability^6,12,13^. Recent single-cell analyses further demonstrate that ageing remodels both epithelial and immune compartments of the colon, establishing a pro-inflammatory tissue environment with altered epithelial-immune crosstalk^13^. Nevertheless, although ageing is known to increase susceptibility to enteric infection, whether this reflects progressive loss of intestinal function or a distinct host response to infection, remains poorly understood.

*Citrobacter rodentium* (CR) is a murine-specific pathogen and is widely used to model enteropathogenic and enterohaemorrhagic *Escherichia coli* (EPEC and EHEC) infections^14,15^. In C57BL/6 mice, CR induces a self-limiting infectious colitis, spreads across 5 distinct phases: establishment, expansion, steady-state, clearance and post clearance tissue recovery. Infection is characterised by colonic crypt hyperplasia (CCH), epithelial barrier disruption, immune cell recruitment, and mucosal cytokine induction^16^. A coordinated epithelial and immune host response includes Il-22 dependent antimicrobial peptide production and epithelial remodelling, neutrophil recruitment, and activation of Th1- and Th17-associated cytokines, including Ifn-γ and Il-17a, which contribute to inflammatory amplification and pathogen clearance^16^.

CR infection is primarily mediated through the injection of type III secretion system (T3SS) effector proteins into intestinal epithelial cells (IECs), where they assemble into a robust and adaptable intracellular signalling network. This effector network remodels epithelial and immune pathways to promote bacterial colonisation and disease^17,18^. Collectively, these effectors target a broad range of host cellular processes, structures, and organelles, including actin cytoskeletal dynamics, NF-κB signalling, cell death pathways, and intracellular trafficking, as well as epithelial tight junctions and mitochondrial function^19^.

We recently showed that deletion of a subset of 12 effectors (Map, NleD1, NleD2, NleH, NleF, NleG1, NleG7, EspJ, EspV, EspK, NleN, and NleK) in a strain named CR_M12_ does not impair colonisation of 8-10 weeks old C57BL/6 mice, although it attenuates epithelial damage and disease outcomes^18^. In contrast, CR_M12_ was fully virulent and resulted in mortality in susceptible C3H/HeN mice, immunocompromised *Il22^-/-^* mice^18^, and mice with a prior history of DSS-colitis^20,21^, indicating that its virulence is strongly influenced by host physiological context. These findings established CR_M12_ as a valuable functional probe to determine how host physiological context shapes the outcome of CR infection.

Here, we show that aged (20-24-month-old) C57BL/6 mice develop exacerbated infectious colitis following CR_WT_ infection compared with young mice (8-10 weeks old). We next asked whether this increased susceptibility reflects complete loss of intestinal function or whether the aged intestine retains the capacity to respond differently to pathogens with distinct virulence repertoires. Using CR_M12_ as a functional probe we demonstrate that although ageing increases susceptibility to wild-type CR (CR_WT_) infection, it does not abolish the adaptive capacity of the intestinal epithelium. Instead, the aged intestine represents a distinct host context that generates divergent epithelial, inflammatory and molecular responses according to pathogen virulence.

## Materials and Methods

### Mouse experiments

All procedures were performed in agreement with the Control and Supervision Rules, 1998 of the Ministry of Environment and Forest Act (Government of India) and the Animal Ethics Committee of the Indian Institute of Science (Approval CAF/Ethics/547/2017). Experiments were performed following principles of Replacement, Reduction, and Refinement (3Rs). Accordingly, to minimise animal use, baseline data for UI and CR_WT_-infected aged mice were used as shared controls for multiple analyses, as indicated in the figure legends. This approach minimized unnecessary duplication of control groups while ensuring robust statistical power and experimental rigour.

All animals were bred and housed in the same vivarium. C57BL/6 mice were housed in a clean air facility, in multiple cages separated based on sex, at 22 ± 2 °C, 55% ± 10% humidity on 12h light/dark cycle. Mice were fed laboratory chow (∼24% protein, 6% oil, and 3% dietary fibres) and water ad libitum. Mice experiments were conducted with 3-5 mice per group. 8-10 weeks-old young mice and 20-24 months-old, aged mice, both male and female were used in all experiments.

### CR infection

Wild type CR (ICC169) and its isogenic mutant CR_M12_^18^ were grown overnight in Lysogeny broth (LB) supplemented with Nalidixic acid (Nal, 50 μg/ml) at 37 °C, 180 rpm, followed by centrifugation (3000 x g) for 10 min and resuspension in sterile 1X PBS. All mice weight were recorded (d0), and mice were infected with 200 μL bacterial suspension in PBS by oral gavage as previously described^22^. Mock-infected mice received 200 μL of sterile PBS. The inocula were used for CFU quantification and was calculated to ∼3 x 10^9^ CFUs. Infections were followed by plating mouse faecal matter at various timepoints on LB + Nal plates as previously described^22^.

### Postmortem analyses

Mice weights were monitored regularly on indicated timepoints. At 6 dpi, postmortem, the colon and caecum were excised, the colon was separated from caecum, weighed using a digital scale, normalised to its length, and recorded. From the distal side of the colon, 0.3 cm was submerged in 4% paraformaldehyde (PFA) and processed for histological analysis, next 2 cm was used for extraction of CR-attached IECs, next 0.5 cm was used for colon explant cultures (described below). To estimate systemic dissemination of CR, liver and spleen were excised, weighed, homogenised in sterile PBS, serially diluted, and plated on LB + Nal plates.

### Histological analyses and immunohistostaining

0.5 cm distal colons were fixed in 4% PFA for 2.5 h, followed by storage in 70% ethanol. The fixed tissues were embedded in paraffin, sectioned at 5 μm, and stained with haematoxylin and eosin (H&E). CCH was quantified in each mouse by measuring >10 well-oriented crypts, and the mean crypt length from each mouse plotted. For immunohistostaining, unstained colon sections were first dewaxed by immersion in Histoclear solution (10 min x 2), followed by immersion in 100% ethanol (10 min x 2), 95% ethanol (3 min x 2), 80% ethanol (3 min), respectively and washes in 1X PBS + 0.1% Tween-20 + 0.1% saponin (PBS-TS) (3 min x 2). The sections were then heated for 20 min in 0.3% trisodium citrate + 0.05% Tween-20 in distilled water (demasking solution, pH=6), cooled, washed in PBS-TS, followed by blocking in PBS-TS + 10% normal donkey serum for 20 min. The slides were incubated overnight with anti-CR polyclonal antibody (1 in 50), anti-Ly6G (1 in 200), Pcna (1 in 500), and/or anti-E-cadherin (1 in 50) antibodies, washed twice (10 min x 2) in PBS-TS, and incubated with appropriate secondary antibodies (1 in 100) and DAPI (1 in 1000, to stain DNA). The slides were washed twice, mounted with ProLong Gold antifade mountant, and imaged and analysed on Zen 2.3 Blue Version (Carl Zeiss Microimaging GmbH, Germany). Pcna^+^ proliferative zone was measured from >10 well-oriented crypts per mouse and plotted as a percentage of respective crypt length. Neutrophil recruitment to the distal colon was quantified by counting the number of neutrophils present in the colonic mucosa, normalised to area, in each mouse. The histological scoring was based on four parameters, crypt hyperplasia, submucosal thickening, immune cell infiltration, and colonic epithelial damage, where scores range from 1-5, wherein 1 is equivalent to a healthy colon. The CR staining and invasion score was determined based on the intensity and distribution of CR staining, the absence or presence of deep crypt localisation, and the extent of bacterial invasion into crypt depth. Scores range from 0-5, where a score of 0 indicates no visible CR staining and an absence of deep crypt localisation.

### Isolation of CR-attached and bystander IECs and proteomic analysis

The experiment was performed thrice, each time with 3-5 aged mice (20-24 months) per group for CR_WT_ and CR_M12_-infected mice and UI controls. In each experiment, IECs were extracted and pooled per group, and further processed (Fig. S3A). As previously described^18^, 2 cm distal colons were excised, sliced open longitudinally, cleaned, and IECs were dissociated from colons by incubating in enterocyte dissociation buffer containing 1X HBSS (Ca and Mg free) + 10 mM HEPES + 1 mM EDTA + 0.5% β-Mercaptoethanol, at 37 °C, at 180 rpm for 45 min. The remaining tissue was discarded, and dissociated IECs were centrifuged (3000 rpm,10 min, RT), washed once with ice-cold PBS (2000 x g for 10 min at 4 °C), and incubated in Mg and Ca-free Dulbecco’s phosphate buffered saline (DPBS) containing 50 μg/ml DNAse I (Thermo Fisher Scientific) (10 min, RT) for DNasing. Next, IECs were passed through 70 μM strainer, centrifuged, washed once with FACS (Fluorescence activated cell sorting) buffer (DPBS + 5% faetal bovine serum [FBS] + 2 mM EDTA), and incubated in 10% Fc Block (Miltenyi Biotec) in FACS buffer for 10 min at 4 °C. Colonic IECS were next stained by anti-CR rabbit polyclonal ab (1 in 1000 dilution) for 20 min at 4 °C, washed twice in filtered MACS (Magnetic activated cell sorting) buffer (DPBS + 2 mM EDTA + 5% BSA), incubated with anti-rabbit magnetic beads in MACS buffer for 20 min at 4 °C, followed by two washes and resuspension in MACS buffer. The dissociated IECs from UI was processed in the same manner till this step and remained resuspended in MACS buffer.

LS column was fitted to magnetic field of MACS separator and preactivated by passing MACS buffer; IEC suspension from CR_WT_/CR_M12_- infected mice was then applied to column and column washed thrice by MACS buffer. The flowthrough (FT) samples were collected and represented predominantly the CR-not associated/bystander (CR^-^) IECs. Magnetically labelled CR^+^ IECs were eluted from column using 5 ml MACS buffer using a magnetic plunger.

MACS elution, MACS FT and UI IECs samples were washed twice in ice-cold PBS, once with FACS buffer, and incubated in FACS buffer containing anti-CR rabbit polyclonal ab (1 in 2000 dilution) for 20 min at 4 °C to relabel CR^+^ IECs. Next, cells were washed, incubated with anti-rabbit PE and anti-Epcam APC antibodies for 30 min at 4 °C, washed twice in sorting buffer (Ca and Mg free DPBS + 1% FBS + 2 mM EDTA), and resuspended in the same. Colonic IECs from UI control (Epcam^+^/CR^-^) was used to gate between CR- and CR^+^ IEC populations (Fig. S3B). Using MACS eluate as input, Epcam^+^/CR^+^ single cells were sorted (∼ [3.1 - 5.3] x 10^5^ for CR_WT_ and ∼ [2 - 4.4] x 10^5^ for CR_M12_), representing CR-attached IECs. From UI control, ∼ [7.3 – 10] x 10^5^ Epcam^+^/CR^-^ IECs were sorted, which was used as the control. Using MACS FT as input, Epcam^+^/CR^-^ single cells were sorted (∼ [2.1 – 9.8] x 10^5^ for CR_WT_ and ∼ [7.1 - 10] x 10^5^ for CR_M12_), representing CR-not-associated or bystander IECs (Fig. S3B). The sorted cells were collected by centrifugation and stored at -70 °C until further processing for proteomic analysis.

### Sample preparation and TMT labelling

Epcam^+^/CR^+^ and Epcam^+^/CR^-^ IECs from CR_WT_ or CR_M12_-infected mice and Epcam^+^/CR^-^ IECs from UI mice (n=3 experiments) were lysed in 100 mM TEAB containing 1% sodium deoxycholate (SDC), 10% isopropanol, 50 mM NaCl, and Protease and Phosphatase inhibitor cocktail (Thermo Fisher Scientific), boiled for 5 min, and re-sonicated. Protein concentration was measured using Quick Start™ Bradford protein assay (BioRad). 15 μg protein were reduced in 5 mM TCEP (1 h), alkylated with 10 mM IAA (30 min), and digested with trypsin (Pierce; 75 ng/μl) (18 h, RT). Peptides were labelled using TMTpro 18-plex label reagent (Thermo Fisher Scientific) according to manufacturer’s instructions. SDC was precipitated with 2% formic acid and removed by centrifugation (10,000 rpm, 5 min), and the supernatant containing TMT-labelled peptides was dried.

### Mass spectrometry analysis

TMT-labelled peptides were fractionated by high-pH-reversed-phase HPLC (Waters XBridge C18, 2.1 x 150 mm, 3.5 μm) on a Dionex ultimate 3000 system, using ammonium hydroxide (0.1%) as mobile phase A and 100% acetonitrile/ 0.1% ammonium hydroxide as mobile phase B. Stepwise separation of peptides was achieved with a gradient elution at 200 μl/min: isocratic for 5 min at 5% phase B, gradient for 40 min to 35% phase B, gradient to 80% phase B in 5 min, isocratic for 5 min, and re-equilibrated to 5% phase B. Fractions were gathered in a 96-well plate every 42 sec, dried, and reconstituted in 50 μl 0.1% formic acid.

Analysis was performed as previously described^18^. Approximately 3 µg of peptides per fraction was analysed on an Orbitrap Ascend mass spectrometer (Dionex Ultimate 3000 system and mass spectrometer [Thermo Fisher Scientific] used for data acquisition) using a Real-Time Search-SPS-MS3 method. Peptides were loaded onto a C18 column (Acclaim PepMap 100, 100 µm × 2 cm, 5 µm, 100 Å) at a flow rate of 10 µl/min and separated over a 120-min low-pH gradient on a nanocapillary reversed-phase column (Acclaim PepMap C18, 75 µm × 50 cm, 2 µm, 100 Å) at 50°C. MS1 scans were collected over a 400-1600 m/z range using an Orbitrap at 120,000 resolution, with standard AGC and auto injection time, and included 2-6 precursor charge states. A dynamic exclusion window of 45 sec was applied with a repeat count of 1, mass tolerance of 10 ppm, and isotope exclusion permitted.

MS2 spectra were acquired in the ion trap at a turbo scan rate with HCD collision energy set to 32% and a maximum injection time of 35 ms. These spectra were explored versus the *Mus musculus* and CR proteomes using the Comet search engine. Static modifications included cysteine carbamidomethylation (+57.0215) and N-terminal/lysine TMTpro (+304.2071), with variable modifications including Asn/Gln deamidation (+0.984) and Met oxidation (+15.9949), allowing up to two variable modifications and a maximum of four peptides per protein. Precursors meeting these parameters were chose for SPS10-MS3 scans, performed at an Orbitrap resolution of 45,000 with normalised HCD collision energy set to 55%, AGC at 200%, and a maximum injection time of 200 ms.

## Proteomics data processing and analysis

Data were normalised across TMT batches by channel sum scaling, with missing value (>60% channel presence) imputed to the minimum batch intensity and further scaled within.

PCA analysis was performed using log_2_ transformed scaled abundances in the Perseus software (version 2.0.11)^23^. Log_2_ fold changes (log_2_FC) were calculated relative to UI controls. Differential expression was assessed using two-sample t-tests (FDR<0.05, or p < 0.05 as mentioned) applying a significance threshold of (|log_2_FC| ≥ 0.5). Heatmaps and hierarchical clustering were generated using Phantasus (v1.25.4)^24^, and volcano plots using VolcaNoseR^25^.

## Faecal water and sodium content analysis

As described previously^18^, to assess faecal water content, freshly collected faecal pellets were placed in pre-weighed 1.5 mL tubes with punctured caps. Tubes containing faeces were weighed and then incubated at 55 °C. The tubes were weighed daily until a constant weight was reached. Wet and dry faecal weights were calculated after subtracting the tube weight and the faecal water content was determined using the following formula:

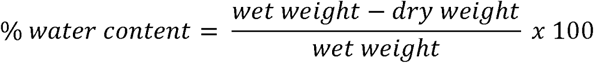

Faecal sodium content was estimated as described earlier^26^. Dry faeces were homogenised in autoclaved double distilled water, centrifuged (100 x g, 1 min), and the supernatant collected in fresh tube. The sodium content of the supernatant was determined using LAQUAtwin metres (Horiba, Japan) and was normalised to the dry weight of the faeces.

## Faecal sample ELISA

Freshly collected faecal pellets were homogenised in 1X PBS containing 0.1% Triton X-100 (1 ml per 100 mg faeces) for 20 min, centrifuged briefly, and supernatant collected and stored at -70 °C. The concentration of Lcn2/NGAL and Mpo in the supernatants were estimated using DuoSet mouse Lipocalin-2 and Mpo ELISA (R&D systems), according to manufacturer’s instructions. Colorimetric readings were obtained using a FLUOstar Omega microplate reader (BMG Labtech). ELISA measurements were performed in duplicates and averaged.

## Cytokine and chemokine profiling from colon explants

0.5 cm distal colon sections were excised, faecal matter removed, tissue weighed, and incubated in sterile RPMI 1640 medium with glutamine (Sigma) + 100 μg/mL streptomycin + 100 U/mL penicillin for 2 h at RT. The tissue was then washed in complete RPMI medium (RPMI 1640 medium with glutamine + 10 mM HEPES, 1 mM sodium pyruvate, 10% FBS, and 100 μg/mL streptomycin + 100 U/mL penicillin) and cultured in the same medium (at 1 mL/0.1 g tissue) for 20 h at 37 °C, 5% CO_2_. The supernatant was extracted, centrifuged (3000 x g, 10 min) to remove cellular debris, and stored at -80 °C until further analysis.

Explant cytokine and chemokine levels from the samples were measured by LEGENDplex kit (BioLegend), according to the manufacturer’s instructions. The panel included Il-22, Il-17a, Il-1β, Ifn-γ, Tnf, Il-18, Cxcl1, Il-6, Il-10, Ccl3, Gm-csf, G-csf, Il-23. Cytokine and chemokine levels were acquired using FACSCalibur flow cytometer (BD Biosciences), and the analysis performed in LEGENDplex data analysis software (BioLegend). Il-23 levels were very low or undetectable in all samples and were excluded from analysis.

## Statistics

Sample size was not predetermined using statistical methods. Experiments were conducted in randomised block structure, including 3-5 mice per group per experiment, with each experiment independently repeated at least twice. The number of mice per experiment is provided in Table S1. Data from independent experiments were pooled for analysis.

Statistical comparisons between two normally distributed groups were performed using a two-tailed t-test; otherwise, a Mann-Whitney test was applied. For comparisons of three or more groups, One-way ANOVA with the indicated post hoc test was used, or, if normality (assessed by Shapiro-Wilk or Kolmogorov-Smirnov tests) was not met, a logarithmic transformation was applied, following by analysis with One-way ANOVA with the indicated post hoc test. Statistical analysis of proteomics data is detailed in the above section. For comparisons of two or more groups across different timepoints, Two-way ANOVA with the indicated post hoc test was used. False discovery rate (FDT, Q=5%) was used to correct for multiple comparisons (Tukey’s post-test), or as implemented by GraphPad Prism (v11.0.2). P values adjusted using FDR, if exceeded 0.05 were considered significant (ns). Data visualisation and statistical analyses were performed using Prism 11.0.2 with details provided in the figure legends.

## Results

### Ageing increases susceptibility to CR infection

To examine the impact of ageing on enteric infection, 20-24-month-old C57BL/6 mice were infected with CR_WT_, with 8-10-week-old mice serving as young controls (Fig. 1A). By 6 days post infection (dpi), aged mice exhibited significant body weight loss, whereas, as previously reported for young mice^16^, they maintained stable body weight (Fig. 1B). Aged mice also displayed increased faecal bacterial shedding, together with higher faecal water content and sodium levels (Fig. 1C-E), indicating a greater bacterial burden accompanied by disruption of fluid-ion homeostasis.

**Fig. 1.**
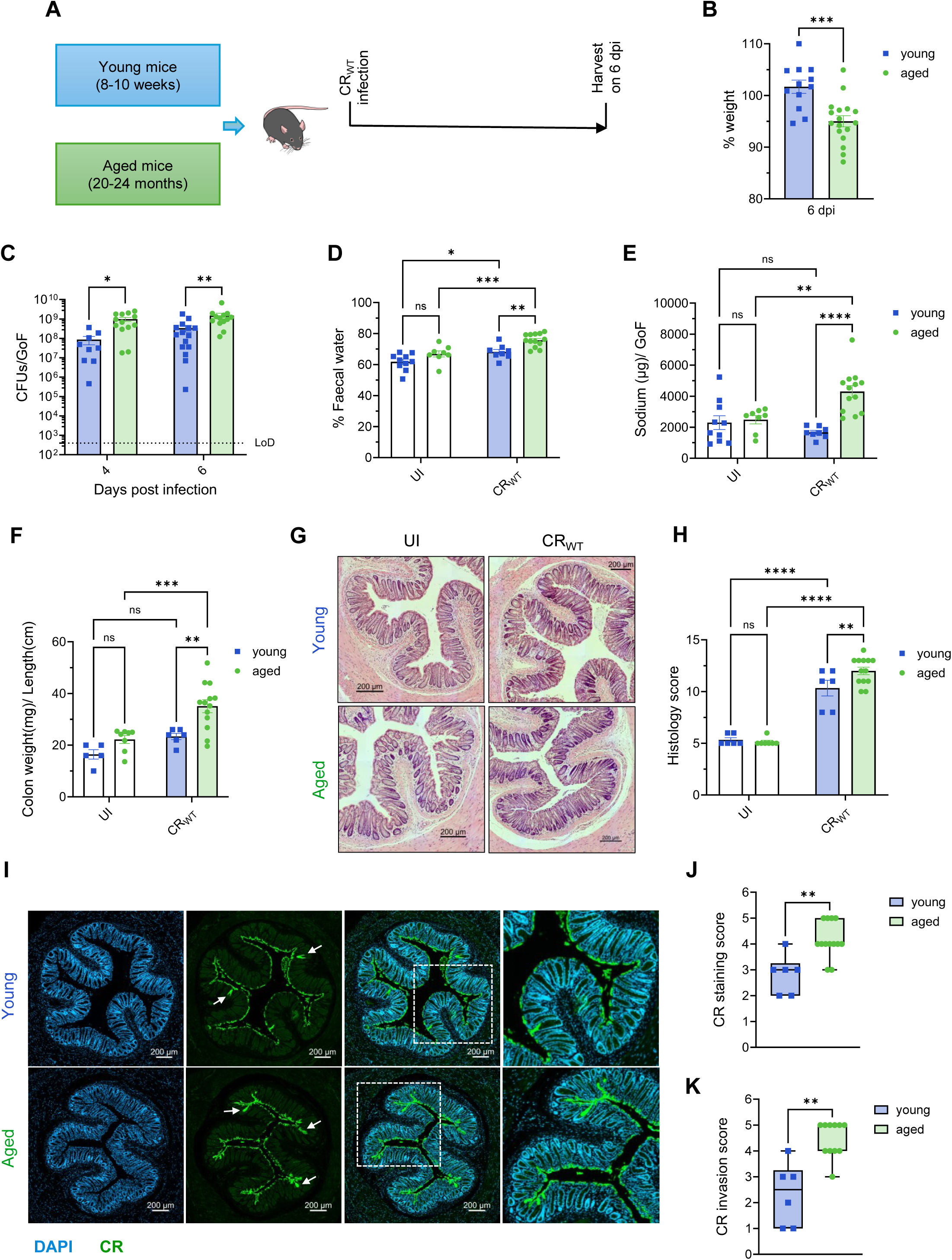
Ageing increases susceptibility to CR infection. (**A**) Schematic representation of the experimental design: 20-24 months-old C57BL/6 mice were infected with CR_WT_. Mice were harvested at 6 dpi. 8-10 weeks-old young mice, infected with CR_WT_ were used as controls. (**B**) Percentage body weight change at 6 dpi. (**C**) Temporal faecal bacterial shedding following CR_WT_ infection. Limit of detection (LoD) is indicated by a dotted black line. (**D**-**E**) Faecal (**D**) water and (**E**) sodium content from CR_WT_-infected mice and UI controls at 6 dpi. (**F**-**K**) Postmortem analysis at 6 dpi. (**F**) Colon weight-to-length ratios, (**G**) representative H&E-stained colon sections. (Scale 200 µm), and (**H**) total histological score. Also see Fig. S1A. (**I**) Representative immunostaining images of colonic sections of mice harvested at 6 dpi. Scale 200 µm. CR (green), and DAPI (blue). (**J**) CR staining and (**K**) CR invasion score representing deep crypt localisation. Each dot represents individual mouse. Data shown are pooled values from at least 2 biological repeats with 3-5 mice per group. Refer to Table S1 for the exact number of mice used for experiments. P values were determined on data plotted as mean ± SEM using two-tailed unpaired Students t-test (**B**), Two-way ANOVA with Sidak’s post-test for multiple comparisons (**C**-**F, H**), and nonparametric Mann-Whitney test (**J**, **K**). ns: not significant; *p < 0.05; **p < 0.01; ***p < 0.001; ****p < 0.0001. GOF: gram of faeces.

Post-mortem analysis at 6 dpi revealed significantly higher colon weight-to-length ratios in aged mice, consistent with increased colonic inflammation^26^ (Fig. 1F). Histological examination of H&E-stained colonic sections demonstrated severe epithelial damage, submucosal thickening, immune cell infiltration and higher histopathology scores in CR_WT_-infected aged mice compared with young controls (Fig. 1G-H, S1A). Despite these differences, both age groups developed comparable CCH and showed a similar expansion of the Pcna-positive proliferative zone relative to their respective uninfected (UI) controls (Fig. 1G and Fig. S1B-D), suggesting that epithelial proliferative responses to infection is preserved during ageing.

Finally, quantification of immunostained colonic sections revealed markedly greater epithelial colonisation and deep crypt localisation of CR in aged mice. In contrast, bacteria in young mice remained predominantly confined to the luminal epithelial surface, a characteristic pattern observed in resistant mouse strains^16,26^ (Fig. 1I-K).

Together, these findings demonstrate that ageing increases susceptibility to CR infection, leading to greater bacterial colonisation and more severe colitis.

### Ageing does not abolish intestinal resilience to CR infection

The increased susceptibility of aged mice to CR infection raised an important question. Does ageing render the intestine functionally equivalent to highly susceptible host settings, or does the aged intestine retain the capacity to distinguish pathogens with different virulence repertoires? To address this, we used the CR mutant CR_M12_ as a functional probe of intestinal resilience. CR_M12_ causes minimal disease in resistant young C57BL/6 mice but causes severe disease in highly susceptible hosts^18^. We therefore reasoned that if ageing resulted in complete loss of intestinal resilience, CR_M12_ would cause disease comparable to CR_WT_ infection.

To test this, aged mice were infected with either CR_WT_ or CR_M12_ (Fig. 2A). Both strains induced a similar reduction in body weight at 6 dpi relative to UI controls (Fig. 2B). However, CR_M12_-infected mice exhibited significantly lower faecal bacterial shedding and reduced colon weight-to-length ratios than CR_WT_-infected mice (Fig. 2C-D), indicating reduced disease severity.

**Fig. 2.**
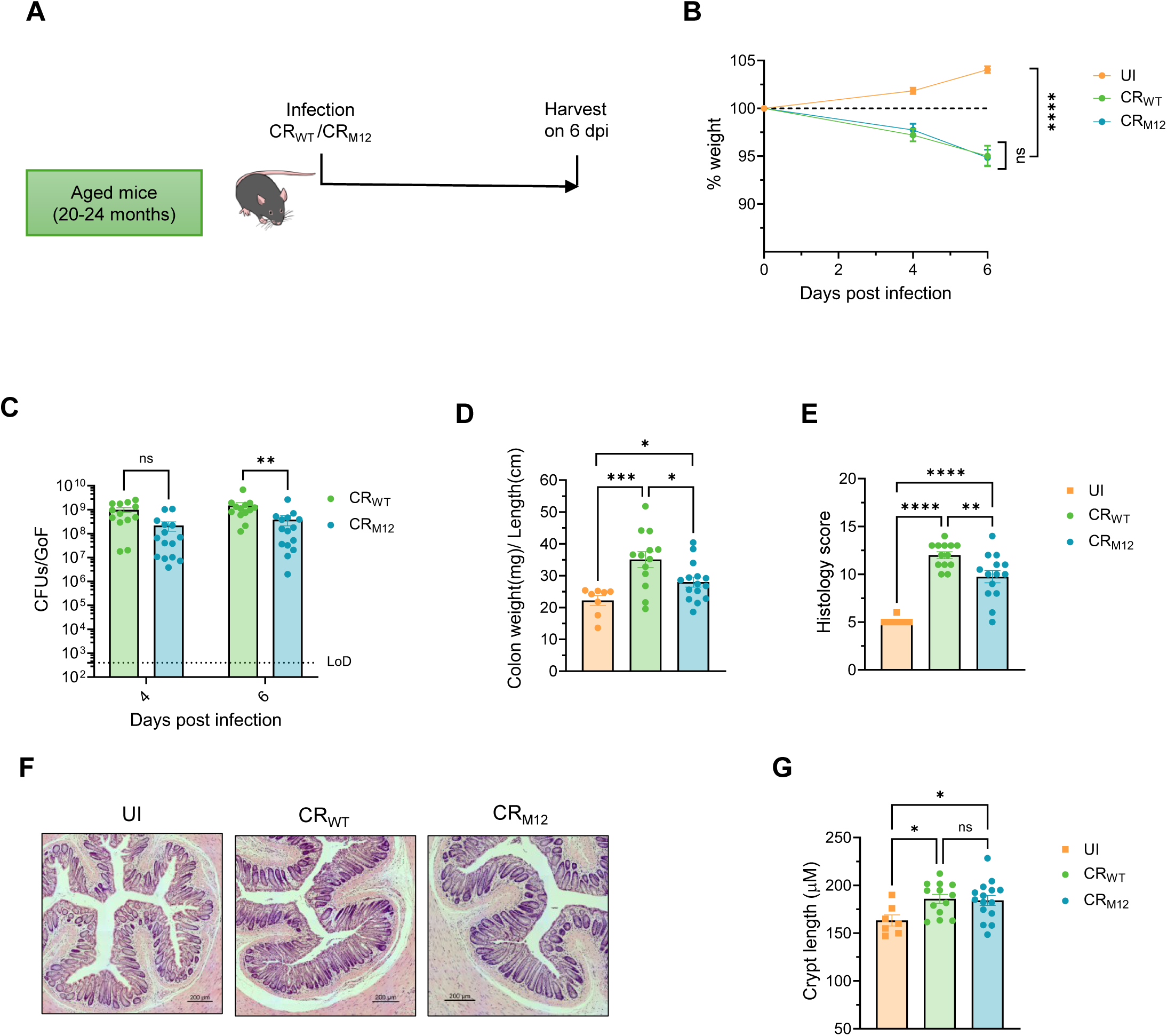
The ageing intestine retains resilience despite increased susceptibility to CR infection. (**A**) Schematic representation of the experimental design: 20-24 months-old C57BL/6 mice were infected with CR_WT_ or isogenic mutant strain CR_M12_. Mice were harvested at 6 dpi. UI mice were used as controls. (**B**-**C**) Temporal (**B**) percentage weight change and (**C**) faecal bacterial shedding following infection. LoD is indicated by a dotted black line. (**D**-**G**) Postmortem analysis at 6 dpi. (**D**) Colon weight-to-length ratios. (**E**) Histological scores for H&E-stained colon sections. Also see Fig. S2A, (**F**) representative H&E-stained distal colon sections of mice harvested at 6 dpi (Scale 200 µm) and (**G**) crypt length measurements. Data for UI and CR_WT_-infected aged mice (**B**-**G**) are same as shown in Fig. 1 and S1B, refer to 3Rs section in Methods. Each dot represents mean (**B**), or individual mouse (**C**-**E**, **G**). Data shown are pooled values from 3 biological repeats with 3-5 mice per group. Refer to Table S1 for the exact number of mice used for experiments. P values were determined on data plotted as mean ± SEM using Two-way ANOVA with Sidak’s post-test for multiple comparisons (**B** and **C**), and One-way ANOVA with Tukey’s post-test for multiple comparisons (**D**, **E**, **G**). ns: not significant; *p < 0.05; **p < 0.01; ***p < 0.001; ****p < 0.0001.

Histological analysis of H&E-stained colonic sections supported these findings. CR_WT_ infection induced marked submucosal thickening, epithelial erosion and inflammatory cell infiltration, whereas CR_M12_ infection resulted in significantly milder pathological changes and lower histopathology scores (Fig. 2E-F). Despite these differences, both CR_WT_- and CR_M12_-infected mice developed comparable CCH (Fig. 2E-G, S2A), demonstrating that epithelial regenerative responses were maintained despite marked differences in disease severity.

Collectively, these data show that ageing does not abolish intestinal resilience. The aged intestine generates distinct responses to pathogens with different virulence repertoires.

### Distinct CR epithelial colonisation is associated with divergent disease outcomes in aged mice

We next asked what distinguishes infections that produce different disease outcomes within the ageing intestine. In highly susceptible hosts, CR_M12_ exhibits epithelial colonisation comparable to CR_WT_^18^. We therefore examined whether differences in epithelial colonisation distinguish CR_WT_ and CR_M12_ infection during ageing by assessing bacterial localisation within the colonic epithelium and quantifying epithelial cells directly associated with bacteria.

Immunohistochemical analysis of colonic sections revealed greater epithelial attachment and deeper crypt localisation of bacteria following CR_WT_ infection. In contrast, CR_M12_ infection was associated with reduced mucosal attachment and limited localisation within the crypt epithelium (Fig. 3A-B).

**Fig. 3.**
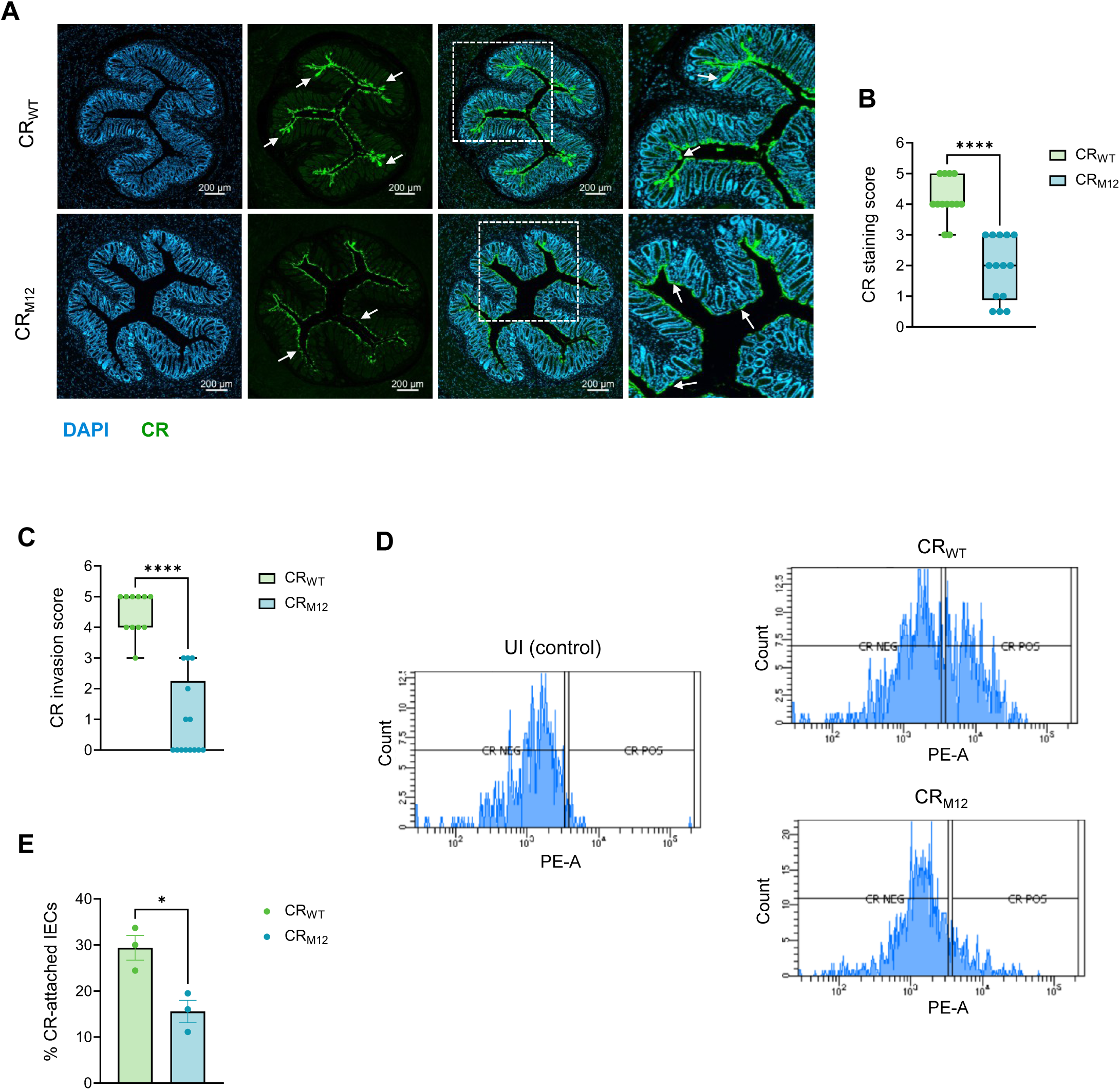
Distinct epithelial colonisation distinguishes divergent disease outcomes during CR infection in aged mice. Aged C57BL/6 mice were infected with CR_WT_ or CR_M12_ and harvested at 6 dpi. UI mice were used as control. (**A**) Representative immunostaining images of colonic sections. Scale 200 µm. CR (green), and DAPI (blue). (**B**) CR staining and (**C**) CR invasion score representing deep crypt localisation. (**D**) Representative histogram of CR-attached IECs (Epcam^+/^CR_WT_^+^ and Epcam^+^/CR_M12_^+^) and (**E**) their quantification as a percentage of total singlet IECs by FACS. CR POS represents Epcam^+^/CR^+^ cells. UI samples were used as a gating control for CR NEG population (Epcam^+^/CR^-^ cells). Data for CR_WT_-infected aged mice (**A**-**C**) are same as shown in Fig. 1, refer to 3Rs section in Methods. Each dot represents independent experimental replicate, a pool of 3-5 mice per group, with experiment performed thrice (**E**), or individual mouse (**B**, **C**). Data shown are pooled values from at least 3 biological repeats with 3-5 mice per group. Refer to Table S1 for the exact number of mice used for experiments. P values were determined on data plotted as mean ± SEM using nonparametric Mann-Whitney test (**B**, **C**, **E**). *p < 0.05; ****p < 0.0001.

To quantify epithelial colonisation more directly, CR-associated epithelial cells were enriched by magnetic-activated cell sorting (MACS), followed by flow cytometric quantification of CR-associated Epcam⁺ IECs (CR⁺ IECs) (Fig. S3A-B). Consistent with the histological findings, ∼30% of Epcam⁺ epithelial cells were CR⁺ following CR_WT_ infection compared with ∼15% following CR_M12_ infection (Fig. 3C-E).

In sum, these observations reveal that the aged intestine was less permissive to epithelial colonisation by CR_M12_ than by CR_WT_.

### The ageing intestine displays preserved epithelial physiology during CR_M12_ infection

Since CR_WT_ and CR_M12_ differ in epithelial colonisation, we next asked whether these differences translate into distinct epithelial physiological responses during ageing. Because epithelial barrier integrity and fluid-ion homeostasis are major determinants of disease severity during CR infection^26^, we examined whether these functions were differentially affected following CR_WT_ and CR_M12_ infection.

To assess epithelial barrier integrity, we first examined E-cadherin localisation by immunohistochemistry. CR_WT_-infected colons exhibited marked disruption of junctional E-cadherin staining, whereas CR_M12_-infected mice largely retained junctional E-cadherin comparable to UI controls (Fig. 4A). Consistent with improved barrier preservation, CR_M12_-infected aged mice also exhibited significantly lower CR burdens in the liver and spleen than CR_WT_-infected mice (Fig. 4B-C). In addition, CR_M12_ infection resulted in significantly lower faecal water and sodium content than CR_WT_ infection (Fig. 4D-E), indicating preservation of epithelial fluid-ion homeostasis.

**Fig. 4.**
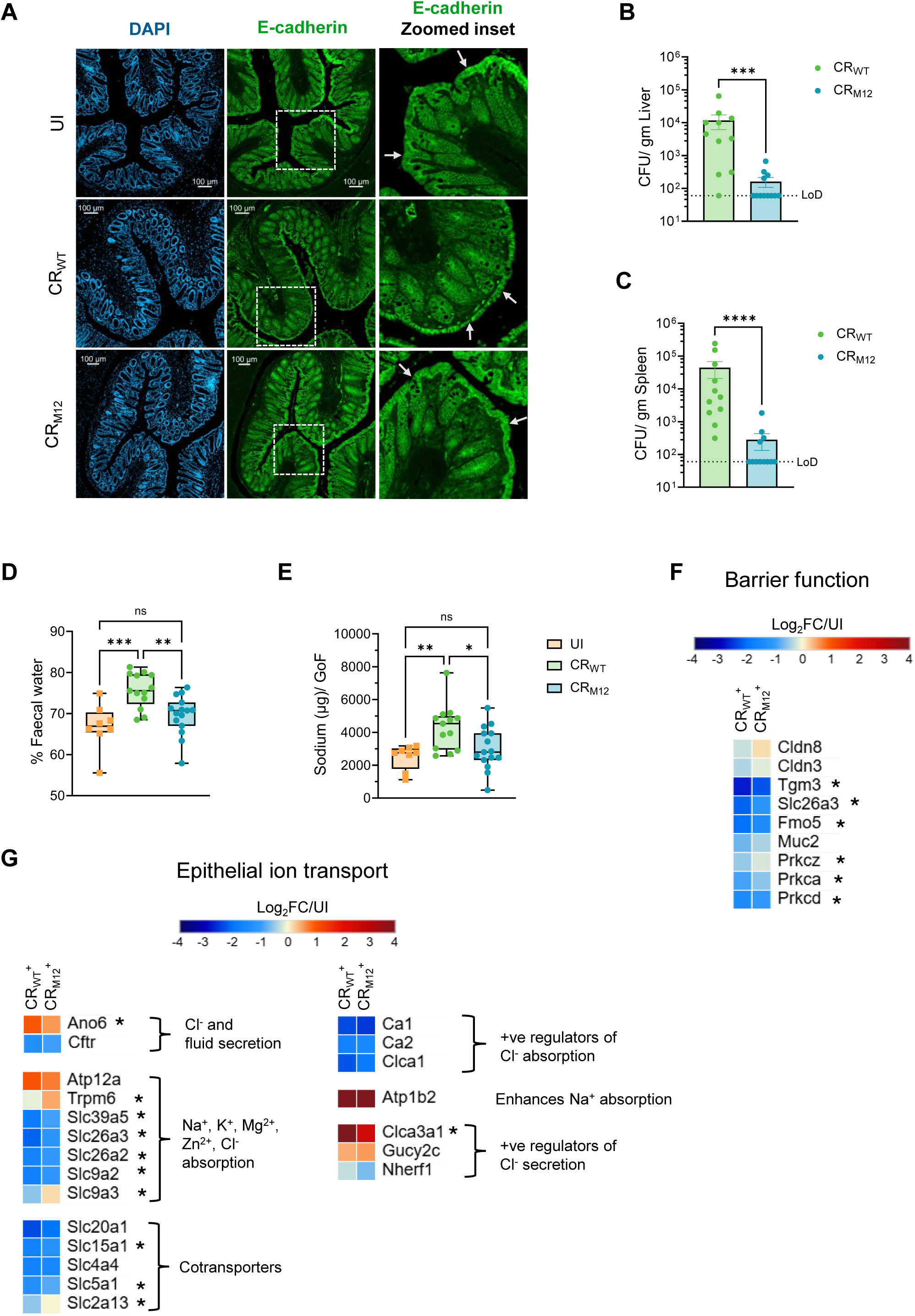
Aged mice display preservation of epithelial barrier function and fluid-ion homeostasis during CR_M12_ infection. Aged C57BL/6 mice were infected with CR_WT_ or CR_M12_ and harvested at 6 dpi. UI mice were used as control. (**A**) Representative immunostaining images of colonic sections of mice harvested at 6 dpi, showing disruption of epithelial barrier. Scale 100 µm. IECs express E- cadherin (green) and DAPI^+^ (blue). The rightmost panel shows zoomed-in E-cadherin inset and white arrows indicate absence or presence of epithelial layer erosion. (**B**-**C**) Extraintestinal dissemination of CR_WT_ and CR_M12_ to (**B**) liver and (**C**) spleen at 6 dpi. LoD is indicated by a dotted black line. (**D**-**E**) Faecal (**D**) water and (**E**) sodium content at 6 dpi. (**F**-**G**) Deep quantitative proteomics analysis of Epcam^+/^CR_WT_^+^ and Epcam^+^/CR_M12_^+^ isolated at 6 dpi using UI samples as control. Heatmap of selected proteins involved in (**F**) barrier function and integrity, and in (**G**) epithelial ion transport. The proteins are further classified into subsets based on function. For **F** and **G**, * refers to proteins significantly different between CR_WT_^+^ and CR_M12_^+^ IECs. Each dot represents individual mouse. Data for UI and CR_WT_-infected aged mice (**D**-**E**) are same as shown in Fig. 1, refer to 3Rs section in Methods. Data shown are pooled values from 3 biological repeats with 3-5 mice per group. Refer to Table S1 for the exact number of mice used for experiments. P values were determined using nonparametric Mann-Whitney test (**B**, **C**) and One-way ANOVA with Tukey’s post-test for multiple comparisons (**D**, **E**). ns: not significant; *p < 0.05; **p < 0.01; ***p < 0.001; ****p < 0.0001. FC: fold changes.

To determine whether these physiological differences were reflected at the molecular level, we analysed barrier- and transport-associated proteins within bacteria-associated (Epcam⁺/CR⁺) IECs identified by quantitative proteomics (Fig. S3A-B). Although core tight junction proteins, including Cldn3 and Cldn8, remained largely unchanged at 6 dpi, both CR_WT_ and CR_M12_ infection reduced the abundance of proteins involved in epithelial barrier maintenance, including Slc26a3, Tgm3, Fmo5 and Prkc isoforms (Prkca and Prkcd) (Fig. 4F). These changes were consistently less pronounced following CR_M12_ infection. Moreover, CR_WT_, but not CR_M12_, reduced the abundance of the atypical protein kinase C isoform Prkcz and the mucus barrier protein Muc2.

Similar differences were observed in epithelial transport programmes. CR_WT_ infection increased the abundance of proteins promoting chloride and fluid secretion while reducing transporters involved in electrolyte absorption and solute transport (Fig. 4G). In contrast, CR_M12_ infection produced substantially milder changes. Key absorptive transporters, including Slc26a3 (Dra), Slc9a3 (Nhe3) and Slc5a1 (Sglt1), were downregulated to lower extent, whereas induction of Ano6 and the chloride secretion regulator Clca3a1 was markedly attenuated compared with CR_WT_ infection (Fig. 4G).

Together, these findings demonstrate preservation of epithelial integrity, fluid-ion homeostasis and epithelial transport programmes in CR_M12_ -infected aged mice.

### The ageing intestine mounts restrained inflammatory responses to CR_M12_ infection

Since CR_M12_-infected aged mice displayed preserved epithelial integrity and fluid-ion homeostasis, we next asked whether these physiological differences were associated with distinct inflammatory responses. We examined neutrophil accumulation, inflammatory mediators and epithelial immune programmes following CR_WT_ and CR_M12_ infection.

Immunohistochemical analysis revealed few Ly6G^high^ neutrophils in UI colons, a marked increase following CR_WT_ infection, and significantly reduced neutrophil accumulation following CR_M12_ infection (Fig. 5A-B). Consistent with these findings, faecal Mpo levels were significantly lower in CR_M12_- than in CR_WT_-infected mice (Fig. 5C), indicating reduced neutrophil accumulation and activity. Similarly, lower faecal Lcn2 levels were observed following CR_M12_ infection (Fig. 5D), consistent with diminished intestinal inflammation.

**Fig. 5.**
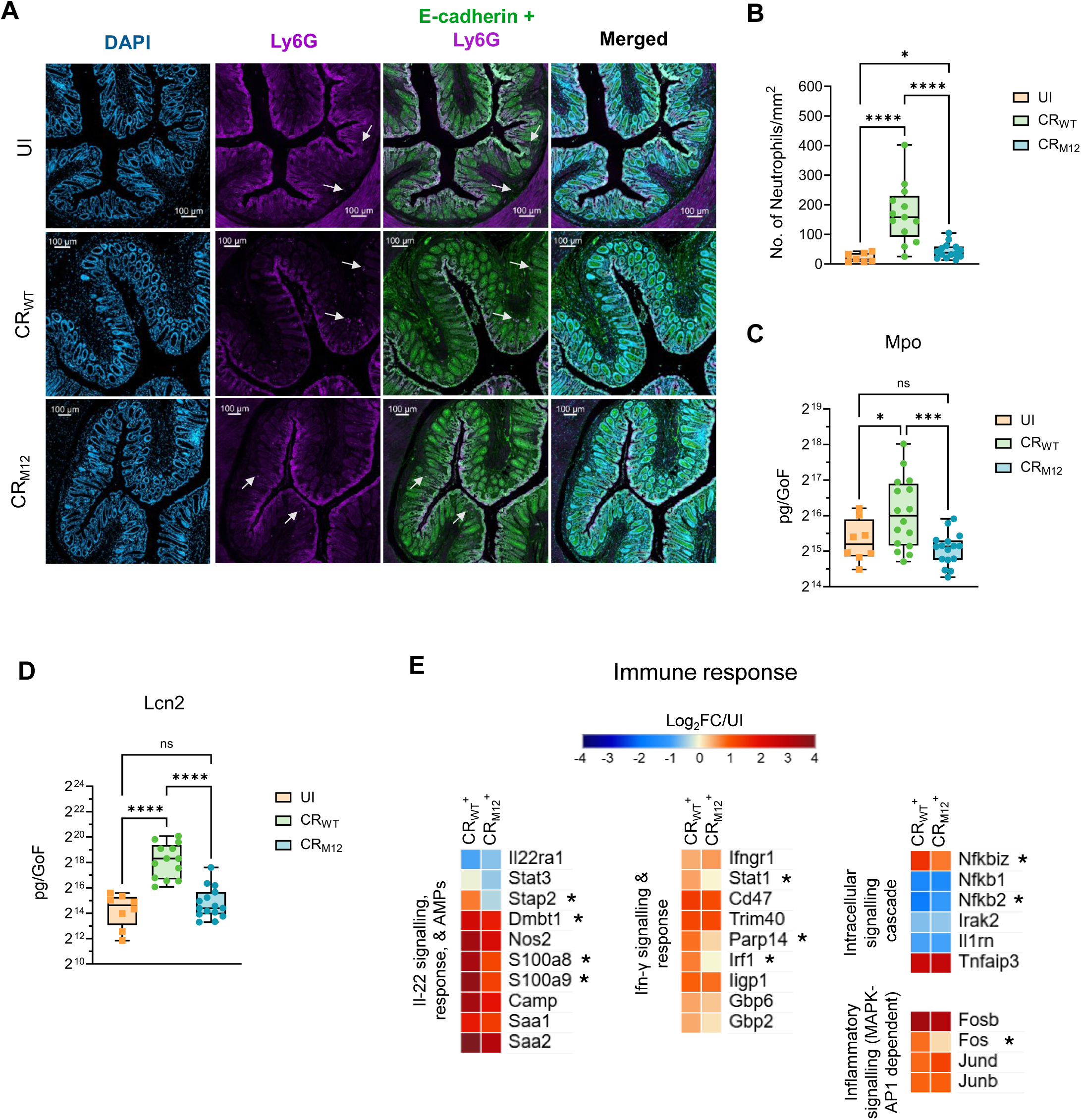
Aged mice display restrained inflammatory responses during CR_M12_ infection. Aged C57BL/6 mice were infected with CR_WT_ or CR_M12_ and harvested at 6 dpi. UI mice were used as control. (**A**) Representative immunostaining images of colonic sections of mice harvested at 6 dpi showing Ly6G^high^ neutrophil recruitment to the colonic mucosa, and (**B**) quantification of the same. Scale 100 µm. IECs express E-cadherin (green) and DAPI^+^ (blue); neutrophils are Ly6G^high^ (violet) and DAPI^+^, but E-cadherin^-^. The rightmost panels show merged images of E-cadherin and Ly6G, and all three channels, respectively. (**C-D**) Faecal (**C**) Mpo and (**D**) Lcn2 levels at 6 dpi. (**E**) Deep quantitative proteomics analysis of Epcam^+/^CR_WT_^+^ and Epcam^+^/CR_M12_^+^ isolated at 6 dpi using UI samples as control. Heatmap of selected proteins involved in host immune response. The proteins are further classified into subsets based on function. * refers to proteins significantly different between CR_WT_^+^ and CR_M12_^+^ IECs. Each dot represents individual mouse. Data shown are pooled values from 3 biological repeats with 3-5 mice per group. Refer to Table S1 for the exact number of mice used for experiments. P values were determined on data plotted as mean ± SEM using One-way ANOVA with Tukey’s post-test for multiple comparisons assuming a lognormal distribution. ns: not significant; *p < 0.05; ***p < 0.001; ****p < 0.0001.

To further characterise the inflammatory response, cytokine production was measured in colonic explant cultures. CR_WT_ infection induced robust production of Ifn-γ, Il-1β and Il-17a, whereas CR_M12_ infection resulted in significantly lower Il-1β levels while the Ifn-γ and Il-17a concentrations were comparable to UI controls (Fig. S4A-C). In contrast, Il-22 and Il-18 were similarly elevated following both CR_WT_ and CR_M12_ infection (Fig. S4D-E). Il-10 and Ccl3 were reduced only following CR_WT_ infection, whereas Tnf, Il-6, Cxcl1, Gm-csf and G-csf remained unchanged in both infection groups (Fig. S4F-L).

Proteomic analysis of Epcam⁺/CR⁺ IECs revealed activation of conserved epithelial immune programmes following both CR_WT_ and CR_M12_ infection, including Il-22-responsive proteins, antimicrobial peptides and Ifn-γ-associated signalling molecules (Fig. 5E). These responses were accompanied by increased abundance of the NF-κB negative regulators Nfkbiz and Tnfaip3, reduced expression of Nfkb1 and Nfkb2, and activation of MAPK–AP-1 signalling, indicating conserved epithelial immune activation during CR infection. However, the magnitude of these responses was consistently lower following CR_M12_ infection, with reduced induction of antimicrobial peptides, Ifn-γ-responsive proteins (Dmbt1, S100a8/S100a9, Stat1 and Irf1) and attenuated modulation of NF-κB-associated signalling proteins (Fig. 5E).

Taken together, these results suggest that preservation of epithelial physiology during ageing is accompanied by restrained inflammatory responses while maintaining core epithelial antimicrobial defence programmes.

### Aged intestinal epithelium presents distinct adaptive molecular programmes in response to CR_WT_ and CR_M12_ infection

The divergent disease outcomes following CR_WT_ and CR_M12_ infection were accompanied by distinct epithelial physiological and inflammatory responses. We therefore asked whether these differences were reflected in the adaptive molecular programmes of the ageing intestinal epithelium. To distinguish epithelial responses resulting from direct bacterial interaction from those occurring more broadly within the tissue, we analysed deep quantitative proteomic of both Epcam⁺/CR⁺ and neighbouring Epcam⁺/CR⁻ bystander IECs (Fig. S3A-B).

Approximately 6,100 proteins were quantified across all samples. Principal component analysis and hierarchical clustering demonstrated clear segregation according to infection status and epithelial cell population, separating UI, Epcam⁺/CR⁺ and Epcam⁺/CR⁻ epithelial cells (Fig. 6A-B). Notably, Epcam⁺/CR_WT_⁺ and Epcam⁺/CR_M12_⁺ epithelial cells clustered more closely with one another than their respective bystander populations, indicating that epithelial responses to direct bacterial contact are broadly conserved, whereas the major molecular differences between CR_WT_ and CR_M12_ infection emerge within the surrounding tissue (Fig. 6B).

**Fig. 6.**
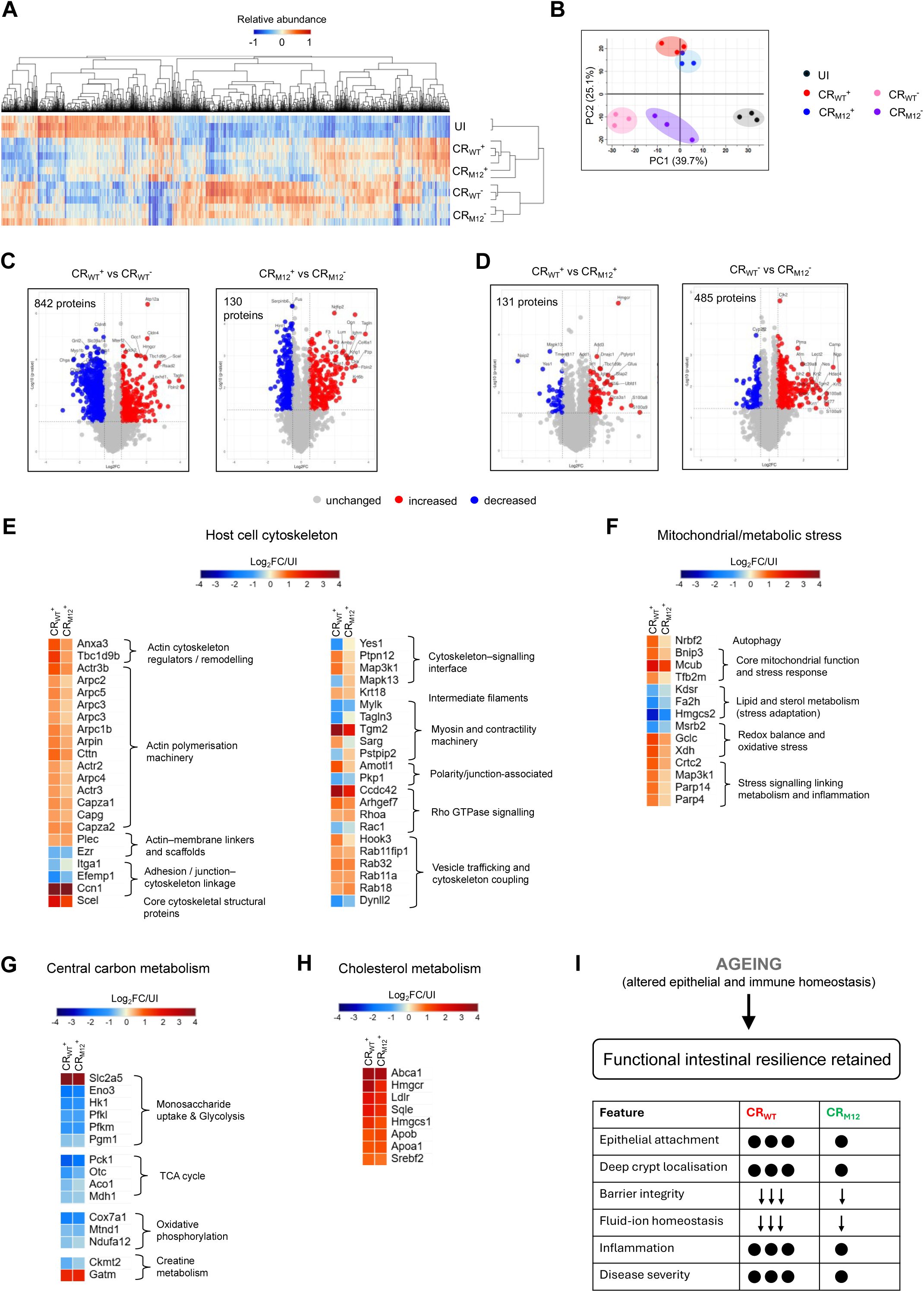
The ageing intestinal epithelium retains distinct adaptive molecular programmes in response to pathogens with different virulence repertoires. Aged C57BL/6 mice were infected with CR_WT_ or CR_M12_ and harvested at 6 dpi. (**A**-**D**) Proteomic analysis was performed on Epcam^+^/CR^+^ and Epcam^+^/CR^-^ IECs from distal colon; Epcam^+^/CR^-^ IECs from UI mice were used as control. (**A**) Hierarchical clustering (one minus cosine similarity) of relative abundance of 3959 proteins that significantly change with infection compared to UI control. (**B**) PCA of mouse proteins detected in CR_WT_^+^, CR_M12_^+^, CR_WT_^-^, CR_M12_^-^, or UI IECs at 6 dpi. Volcano plots of significantly (y-axis) and differentially (x-axis) regulated proteins upon CR_WT_ or CR_M12_ infection relative to UI control, (**C**) CR_WT_^+^ vs CR_WT_^-^, and CR_M12_^+^ vs CR_M12_^-^, (**D**) CR_WT_^+^ vs CR_M12_^+^, and CR_WT_^-^ vs CR_M12_^-^. (**E**-**H**) Heatmaps of selected proteins involved in (**E**) cell cytoskeleton, (**F**) mitochondrial and metabolic stress, (**G**) central carbon and (**H**) cholesterol metabolism following CR_WT_ and CR_M12_ infection of aged mice at 6 dpi. Data shown are pooled values from at least 3 biological repeats with 3-5 mice per group. (I) Proposed working model illustrating preserved functional intestinal resilience during ageing. Although ageing increases susceptibility to CR infection, the ageing intestine retains the capacity to distinguish pathogens with different virulence repertoires. Compared with CR_WT_, CR_M12_ exhibits reduced epithelial colonisation, preservation of epithelial physiology, restrained inflammatory responses and a distinct pattern of epithelial molecular remodelling, resulting in attenuated disease.

Comparison of CR-associated and bystander epithelial cells revealed striking differences in the extent of epithelial remodelling between CR_WT_ and CR_M12_ infection. During CR_WT_ infection, 842 proteins were differentially regulated between CR⁺ and CR⁻ epithelial cells, whereas only 130 proteins differed following CR_M12_ infection (Fig. 6C; Table S2), indicating substantially greater epithelial remodelling in response to CR_WT_. Most of these differences reflected proteins regulated in opposite directions between CR-associated and bystander epithelial cells (728 of 842 proteins during CR_WT_ infection and 122 of 130 proteins during CR_M12_ infection), whereas comparatively few proteins changed in the same direction (114 and 8 proteins, respectively) (Table S2). These findings indicate that epithelial cells directly interacting with bacteria and neighbouring bystander cells adopt fundamentally distinct molecular responses rather than representing a continuum of the same programme. Although CR_WT_- and CR_M12_-associated epithelial cells shared broadly similar proteomic profiles, their corresponding bystander epithelial populations diverged considerably (Fig. 6D), suggesting that pathogen virulence drives extensive tissue-wide epithelial remodelling beyond the subset of epithelial cells directly colonised by CR. These findings indicate that pathogen virulence is associated with epithelial remodelling beyond cells directly associated with bacteria, revealing a tissue-wide component of the aged epithelial response.

Distinct adaptive programmes also emerged within CR-associated epithelial cells. CR_WT_ infection induced extensive remodelling of pathways involved in cytoskeletal organisation, epithelial architecture, mitochondrial function, central carbon metabolism, lipid metabolism, redox homeostasis, proteostasis, cell death and cellular stress responses, together with coordinated regulation of transcriptional, epigenetic and vesicle trafficking programmes (Fig. 6E-H, S5A; Tables S3-S4). Although many of these programmes were also induced following CR_M12_ infection, they occurred with consistently lower magnitude and complexity. These findings demonstrate that pathogens expressing distinct virulence profiles drive quantitatively distinct adaptive programmes within the ageing intestinal epithelium, spanning epithelial architecture, cellular metabolism and tissue-wide remodelling.

Together, these findings demonstrate that although ageing increases susceptibility to CR infection, it does not eliminate the adaptive capacity of the intestinal epithelium (Fig. 6I). The ageing intestine retains distinct molecular programmes that underlie differential epithelial and inflammatory responses, ultimately leading to divergent disease outcomes following infection with pathogens carrying different virulence repertoires.

## Discussion

Ageing is widely associated with increased susceptibility to enteric infection and is often viewed as a state of progressive intestinal dysfunction^27–29^. Here, using CR to model infection in aged mice, our findings support the former but challenge the latter. Although aged mice developed more severe disease following CR_WT_ infection, the aged intestine retained the capacity to distinguish pathogens with different virulence repertoire and generated correspondingly distinct epithelial, inflammatory and molecular responses.

CR_WT_ infection in aged mice resulted in hallmark features of severe infectious colitis^26^, including increased epithelial colonisation, barrier disruption, inflammatory activation and extensive epithelial remodelling, resulting in substantially greater disease than observed in young mice. In contrast, infection with the CR_M12_ strain produced milder epithelial and inflammatory perturbations, characterised by reduced epithelial attachment and deep crypt localisation, preservation of epithelial physiology, restrained inflammatory responses and attenuated molecular remodelling. Importantly, CR_M12_ has previously been shown to cause severe disease and mortality in susceptible hosts, including Il-22-deficient mice, C3H/HeN mice and mice with a history of DSS-induced colitis^18,20^. The finding that CR_M12_ caused lower disease in aged mice therefore indicates that increased susceptibility during ageing does not represent complete loss of intestinal function. Rather, ageing lowers the threshold for severe disease while preserving the capacity of the intestine to generate distinct responses to pathogens encoding different effector profiles.

Although CR_M12_ infection in aged mice retained many of the characteristics previously described in young C57BL/6 mice^18^, including reduced epithelial pathology, preservation of barrier function and attenuated inflammatory responses relative to CR_WT_ infection, several notable differences emerged. Unlike young mice, in which CR_M12_ achieves WT levels of intestinal colonisation while inducing reduced Il-22 production and CCH^18^, aged mice exhibited significantly lower epithelial colonisation and bacterial shedding and showed similar levels of Il-22, Il-18 and epithelial proliferative responses compared to CR_WT_ infection. These observations indicate that ageing modifies the relationship between bacterial colonisation, epithelial physiology and inflammatory responses during CR infection, suggesting that the biological consequences of pathogen virulence are influenced by the aged intestinal environment.

Proteomic analysis showed that the molecular divergence occurred within bystander epithelial cells that are not directly associated with CR. Although CR-associated epithelial cells displayed broadly similar global molecular responses following CR_WT_ and CR_M12_ infection, neighbouring bystander epithelial cells exhibited substantially greater divergence, indicating that pathogen virulence shapes epithelial adaptation at the tissue level rather than solely within cells in direct contact with bacteria. This observation suggests that epithelial responses to enteric infection involve coordinated communication across the intestinal epithelium and raises the possibility that tissue-wide epithelial adaptation contributes to determining infection outcome. Defining the mechanisms underlying this coordinated epithelial response represents an important direction for future investigation.

Our findings place ageing within a broader framework of host responses to enteric infections. Previous studies have established that ageing increases susceptibility to pathogens such as *Salmonella*, *Listeria*, *Campylobacter* and *Clostridioides difficile*, largely through impaired immune responses, microbiota dysbiosis and reduced colonisation resistance^8,11,30,31^. Our findings extend these observations by demonstrating that susceptibility should not be interpreted as complete intestinal dysfunction. Instead, the aged intestine retains the capacity to mount distinct epithelial, inflammatory and molecular responses to pathogens with distinct virulence repertoires. This distinction provides a more nuanced view of intestinal ageing, in which increased age-related disease severity reflects altered host responsiveness rather than uniform functional collapse.

Collectively, our work supports a model in which ageing shifts the balance between susceptibility and adaptation rather than producing a uniformly dysfunctional intestine. The aged epithelium remains capable of mounting coordinated defence, repair and metabolic responses, yet these adaptive programmes become insufficient to fully compensate for infection with highly virulent pathogens. More broadly, these findings suggest that preserving epithelial adaptive responses, rather than simply augmenting antimicrobial immunity, may represent an important strategy for maintaining intestinal health during ageing.

## Data Availability

All materials, key resources, and raw data underlying each result are available on Figshare, as detailed in the files “*Reagents and Resources Table*” and “*Raw Data*”. Figshare link is as follows: https://doi.org/10.6084/m9.figshare.32837150

Proteomics data have been deposited in the Proteome Xchange consortium via the PRIDE repository under accession number PXD079234.

Reviewer access details

Project accession: PXD079234

Token: StUTbMReWNmG

## Supplementary Figures

**Supplementary Figure 1. Histological analysis and immunohistochemistry following CR infection in aged mice** (**A**-**D**) Aged C57BL/6 mice were infected with CR_WT_ and harvested at 6 dpi. Infected young mice were used as control. (**A**) Histological scoring based on epithelial damage, submucosal thickening, immune cell infiltration and hyperplasia. (**B**) Crypt length measurements. (**C**) Representative immunostaining images of colonic sections of mice harvested at 6 dpi. Scale 100 µm. DAPI (blue), and Pcna (violet). (**D**) Quantification of Pcna staining as a percentage of total crypt length. Each dot represents individual mouse. Data shown are pooled values from at least 2 biological repeats with 3-5 mice per group. Refer to Table S1 for the exact number of mice used for experiments. P values were determined on data plotted as mean ± SEM using Two-way ANOVA with Sidak’s post-test for multiple comparisons. ns: not significant; *p < 0.05; **p < 0.01; ****p < 0.0001.

**Supplementary Figure 2. Histology scoring following CR_M12_ infection in aged mice** Aged C57BL/6 mice were infected with CR_WT_ or CR_M12_ and harvested at 6 dpi. UI aged mice were used as control. (**A**) Histological scoring based on epithelial damage, submucosal thickening, immune cell infiltration and hyperplasia. Data for UI and CR_WT_-infected aged mice are same as shown in Figs. S1A, refer to 3Rs section in Methods. Data shown are pooled values from 3 biological repeats with 3-5 mice per group. Refer to Table S1 for the exact number of mice used for experiments.

**Supplementary Figure 3.** Schematics of isolation of Epcam^+^/CR^+^ and Epcam^+^/CR^-^ IECs from infected mice (A) Schematic representation of sample preparation using MACS followed by FACS leading to deep quantitative proteomic analysis of CR-attached/ infected IECs (Epcam^+^/CR^+^) and CR-not associated/bystander (Epcam^+^/CR^-^) IECs at 6 dpi with CR_WT_ or CR_M12_. (**B**) Detailed schematic representation of FACS sorting (mentioned in **A**) of MACS Elution (enriched in CR^+^ IECs) and MACS FT fraction (containing CR^-^ IECs) to isolate Epcam^+^/CR^+^ and Epcam^+^/CR^-^ IECs. Samples for UI mice were used as a negative control to gate CR^-^ population.

**Supplementary Figure 4. Cytokines and chemokines upon CR_M12_ infection of aged mice** Aged C57BL/6 mice were infected with CR_WT_ or CR_M12_ and harvested at 6 dpi. UI mice were used as control. Cytokine and chemokine analysis of colonic explant cultures at 6 dpi for (**A**) Ifn-γ, (**B**) Il-1β, (**C**) Il-17a, (**D**) Il-22, (**E**) Il-18, (**F**) Il-10, (**G**) Tnf, (**H**) Il-6, (**I**) Cxcl1, (**J**) Ccl3, (**K**) Gm-csf, and (**L**) G-csf. Each dot represents individual mouse. Data shown are pooled values from 3 biological repeats with 3-5 mice per group. Refer to Table S1 for the exact number of mice used for experiments. P values were determined on data plotted as mean ± SEM using One-way ANOVA with Tukey’s post-test for multiple comparisons on lognormal value. ns: not significant; *p < 0.05; **p < 0.01; ****p < 0.0001.

**Supplementary Figure 5.** Altered cell death pathway regulation in CR_WT_^+^ and CR_M12_^+^ IECs in aged mice (A) Heatmap of selected proteins involved in cell death pathways. Data shown are pooled values from at least 3 biological repeats with 3-5 mice per group.

## References

1 DeJong, E. N., Surette, M. G. & Bowdish, D. M. E. The Gut Microbiota and Unhealthy Aging: Disentangling Cause from Consequence. Cell Host Microbe 28, 180–189 (2020). 10.1016/j.chom.2020.07.013

2 Kim, H. H. & Dixit, V. D. Metabolic regulation of immunological aging. Nat Aging 5, 1425–1440 (2025). 10.1038/s43587-025-00921-2

3 Li, X. et al. Inflammation and aging: signaling pathways and intervention therapies. Signal Transduct Target Ther 8, 239 (2023). 10.1038/s41392-023-01502-8

4 Nalapareddy, K., Zheng, Y. & Geiger, H. Aging of intestinal stem cells. Stem Cell Reports 17, 734–740 (2022). 10.1016/j.stemcr.2022.02.003

5 Pentinmikko, N. et al. Notum produced by Paneth cells attenuates regeneration of aged intestinal epithelium. Nature 571, 398–402 (2019). 10.1038/s41586-019-1383-0

6 Gamez-Macias, P. E. et al. Intestinal Permeability, Gut Inflammation, and Gut Immune System Response Are Linked to Aging-Related Changes in Gut Microbiota Composition: A Study in Female Mice. J Gerontol A Biol Sci Med Sci 79 (2024). 10.1093/gerona/glae045

7 Perkins, E. H., Makinodan, T. & Seibert, C. Model approach to immunological rejuvenation of the aged. Infect Immun 6, 518–524 (1972). 10.1128/iai.6.4.518-524.1972

8 Zangoui, P., Singh, E., Sheetz, M. P. & Kenney, L. J. Low-frequency ultrasound treatment reduces susceptibility to Salmonella infection in aged mice. J Bacteriol 207, e0017625 (2025). 10.1128/jb.00176-25

9 Han, S. N. et al. Vitamin E supplementation increases T helper 1 cytokine production in old mice infected with influenza virus. Immunology 100, 487–493 (2000). 10.1046/j.1365-2567.2000.00070.x

10 Murciano, C. et al. Impaired immune response to Candida albicans in aged mice. J Med Microbiol 55, 1649–1656 (2006). 10.1099/jmm.0.46740-0

11 Ren, Z. et al. Effect of age on susceptibility to Salmonella Typhimurium infection in C57BL/6 mice. J Med Microbiol 58, 1559–1567 (2009). 10.1099/jmm.0.013250-0

12 Liu, A. et al. Aging Increases the Severity of Colitis and the Related Changes to the Gut Barrier and Gut Microbiota in Humans and Mice. J Gerontol A Biol Sci Med Sci 75, 1284–1292 (2020). 10.1093/gerona/glz263

13 Sirvinskas, D. et al. Single-cell atlas of the aging mouse colon. iScience 25, 104202 (2022). 10.1016/j.isci.2022.104202

14 Mundy, R., MacDonald, T. T., Dougan, G., Frankel, G. & Wiles, S. Citrobacter rodentium of mice and man. Cell Microbiol 7, 1697–1706 (2005). 10.1111/j.1462-5822.2005.00625.x

15 Jordan, S., Frankel, G. & Mishra, V. Citrobacter rodentium. Trends Microbiol 33, 1132–1133 (2025). 10.1016/j.tim.2025.02.018

16 Mullineaux-Sanders, C. et al. Citrobacter rodentium-host-microbiota interactions: immunity, bioenergetics and metabolism. Nat Rev Microbiol 17, 701–715 (2019). 10.1038/s41579-019-0252-z

17 Ruano-Gallego, D. et al. Type III secretion system effectors form robust and flexible intracellular virulence networks. Science 371 (2021). 10.1126/science.abc9531

18 Biswas, P. et al. The accessory type III secretion system effectors collectively shape intestinal inflammatory infection outcomes. Gut Microbes 17, 2526134 (2025). 10.1080/19490976.2025.2526134

19 Sanchez-Garrido, J., Ruano-Gallego, D., Choudhary, J. S. & Frankel, G. The type III secretion system effector network hypothesis. Trends Microbiol 30, 524–533 (2022). 10.1016/j.tim.2021.10.007

20 Biswas, P., Mishra, V., Sanchez-Garrido, J. & Frankel, G. Context-dependent epithelial and immune programs shape intestinal resilience or vulnerability following prior colitis. Cell Mol Gastroenterol Hepatol, 101826 (2026). 10.1016/j.jcmgh.2026.101826

21 Sugihara, K. & Kamada, N. Protection or Vulnerability? The Context-dependent memory of intestinal inflammation. Cellular and Molecular Gastroenterology and Hepatology (2026).

22 Crepin, V. F., Collins, J. W., Habibzay, M. & Frankel, G. Citrobacter rodentium mouse model of bacterial infection. Nat Protoc 11, 1851–1876 (2016). 10.1038/nprot.2016.100

23 Tyanova, S. et al. The Perseus computational platform for comprehensive analysis of (prote)omics data. Nat Methods 13, 731–740 (2016). 10.1038/nmeth.3901

24 Kleverov, M. et al. Phantasus, a web application for visual and interactive gene expression analysis. Elife 13 (2024). 10.7554/eLife.85722

25 Goedhart, J. & Luijsterburg, M. S. VolcaNoseR is a web app for creating, exploring, labeling and sharing volcano plots. Sci Rep 10, 20560 (2020). 10.1038/s41598-020-76603-3

26 Mishra, V. et al. Rehydration rescues Il22(-/-) mice from lethal Citrobacter rodentium infection. Nat Commun 17, 306 (2025). 10.1038/s41467-025-67006-x

27 Soenen, S., Rayner, C. K., Jones, K. L. & Horowitz, M. The ageing gastrointestinal tract. Curr Opin Clin Nutr Metab Care 19, 12–18 (2016). 10.1097/MCO.0000000000000238

28 Wang, Y. et al. Trends in intestinal aging: From underlying mechanisms to therapeutic strategies. Acta Pharm Sin B 15, 3372–3403 (2025). 10.1016/j.apsb.2025.05.011

29 Shen, X., Gao, X. & Wang, L. Intestinal aging-related immune dysfunction: mechanisms and interventions. Acta Biochim Biophys Sin (Shanghai) 58, 183–200 (2026). 10.3724/abbs.2025157

30 Alam, M. S., Gangiredla, J., Hasan, N. A., Barnaba, T. & Tartera, C. Aging-Induced Dysbiosis of Gut Microbiota as a Risk Factor for Increased Listeria monocytogenes Infection. Front Immunol 12, 672353 (2021). 10.3389/fimmu.2021.672353

31 Peniche, A. G., Spinler, J. K., Boonma, P., Savidge, T. C. & Dann, S. M. Aging impairs protective host defenses against Clostridioides (Clostridium) difficile infection in mice by suppressing neutrophil and IL-22 mediated immunity. Anaerobe 54, 83–91 (2018). 10.1016/j.anaerobe.2018.07.011

