## Supplementary Figures for "*Citrobacter rodentium* infection reveals preserved intestinal resilience in aged intestine"

### Supplementary Figure 1

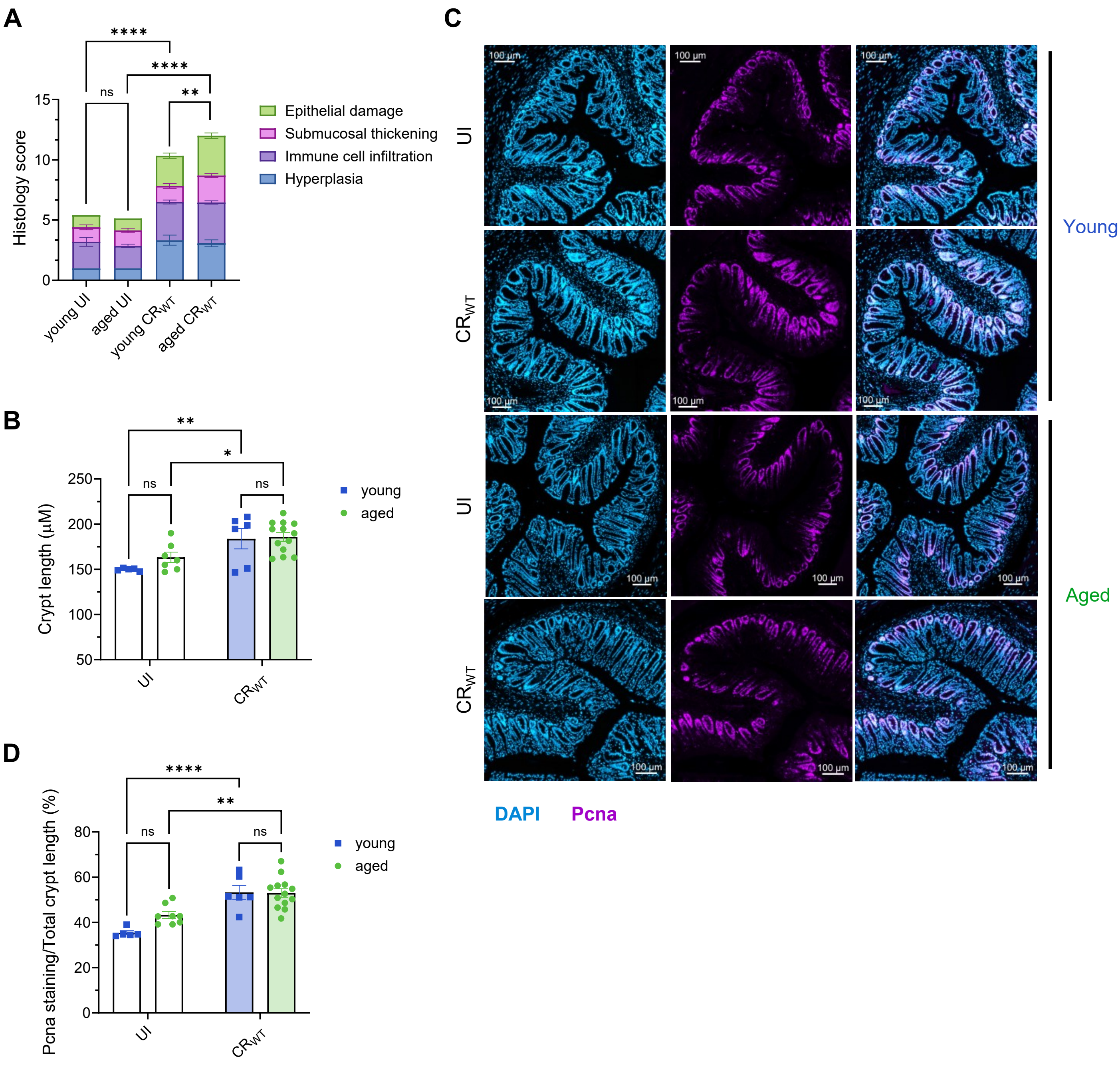

### Supplementary Figure 2

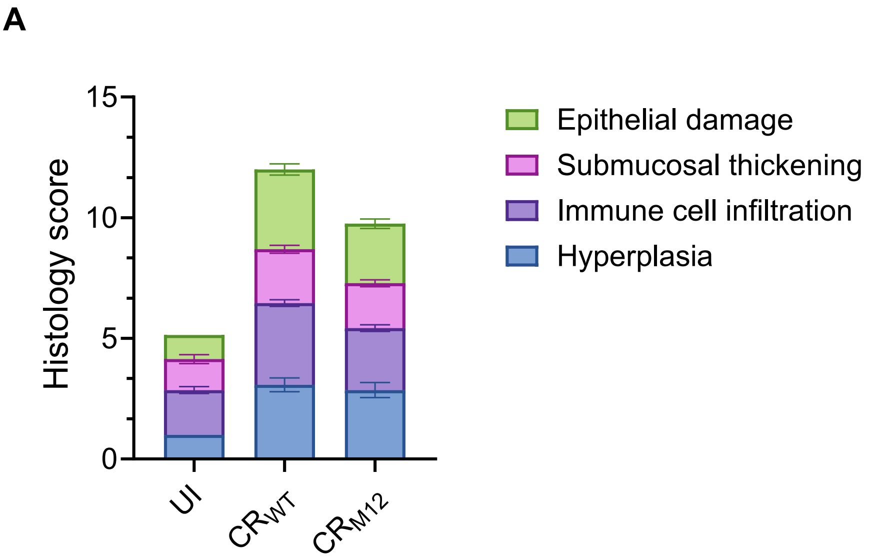

### Supplementary Figure 3

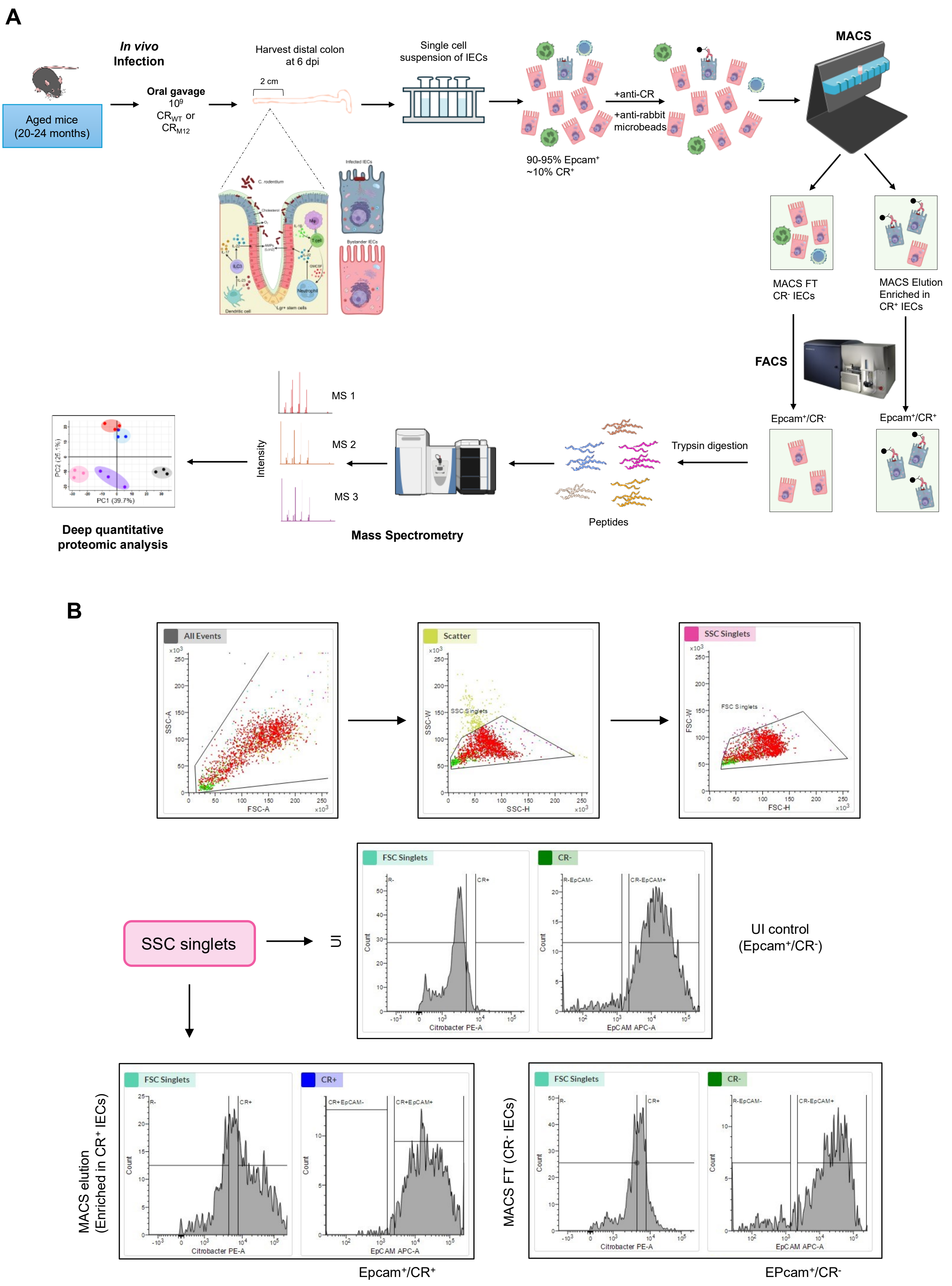

### Supplementary Figure 4

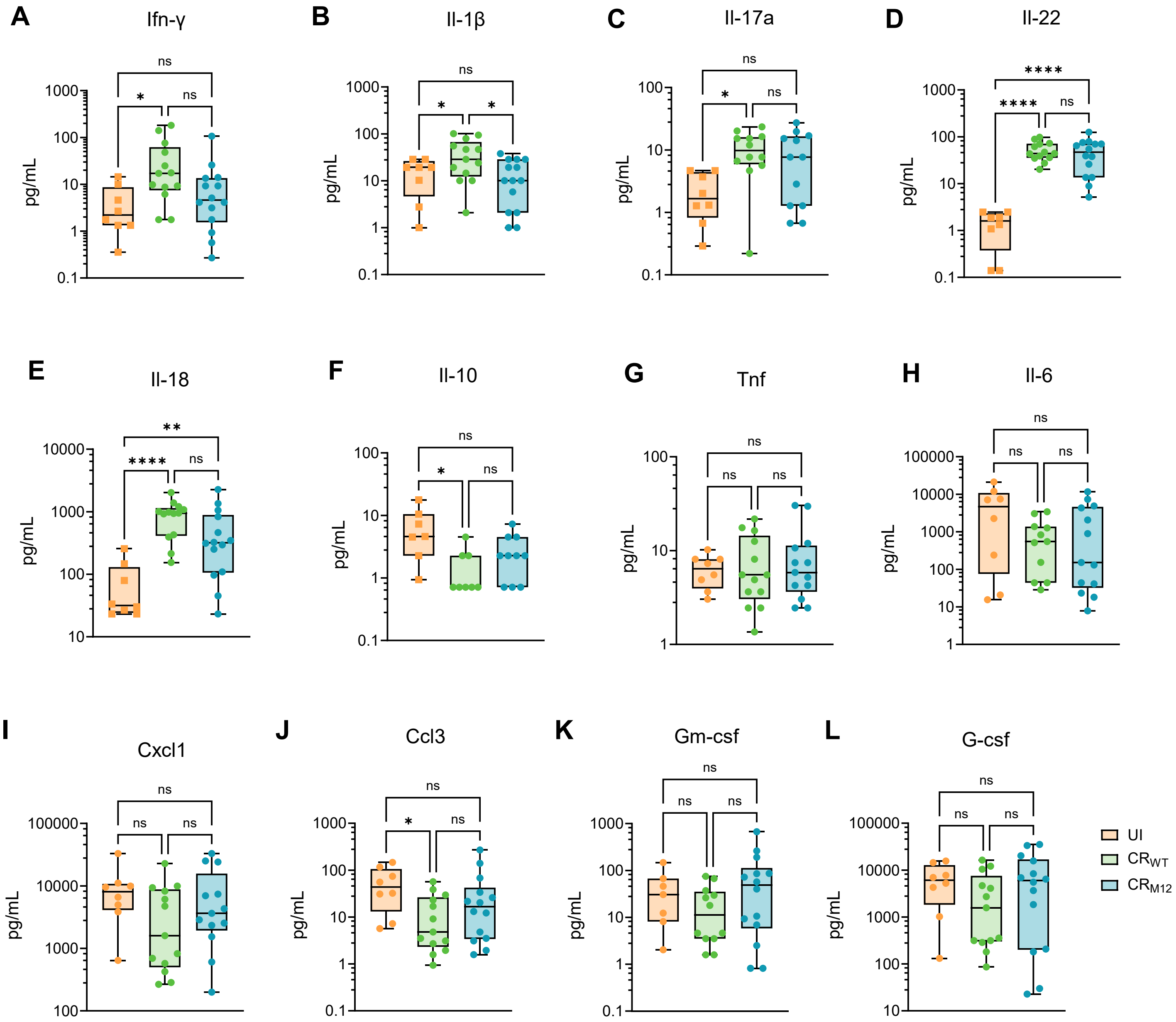

### Supplementary Figure 5

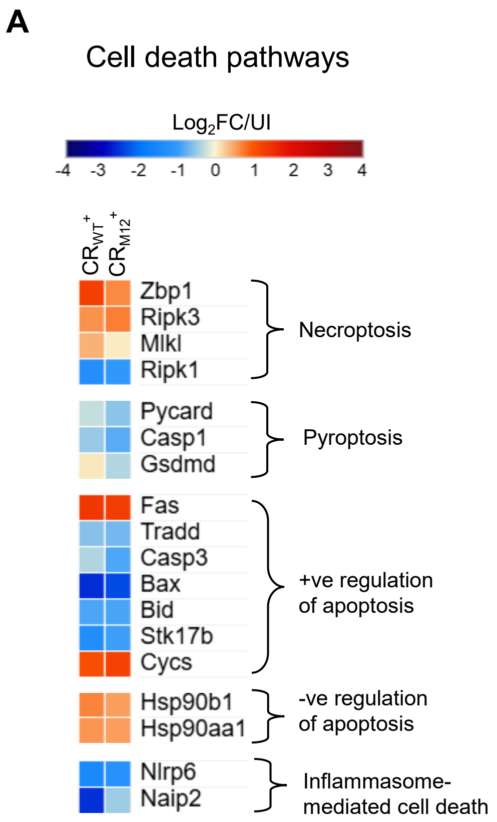

**Supplementary Table 1.** Mice numbers

| Figure | | Samples | Young | Aged | | |
|---|---|---|---|---|---|---|
| 1 | B | Mice no. | 12 | 17 | | |
| | C | | 13 | 15 | | |
| | | | UI young | $CR_{WT}$ young | UI aged | $CR_{WT}$ aged |
| | D | Mice no. | 10 | 8 | 8 | 13 |
| | E | | 10 | 8 | 8 | 13 |
| | F | | 5 | 8 | 6 | 13 |
| | | Samples | UI young | $CR_{WT}$ young | UI aged | $CR_{WT}$ aged |
| | G | Mice no. | 6 | 6 | 7 | 13 |
| | H | | 6 | 6 | 7 | 13 |
| | | Samples | Young | Aged | | |
| | I | Mice no. | 6 | 13 | | |
| | J | | 6 | 13 | | |
| | K | | 6 | 11 | | |
| 2 | | Samples | UI | $CR_{WT}$ | $CR_{M12}$ | |
| | B | Mice no. | 7 | 17 | 18 | |
| | | Samples | $CR_{WT}$ | $CR_{M12}$ | | |
| | C | Mice no. | 13 | 15 | | |
| | | Samples | UI | $CR_{WT}$ | $CR_{M12}$ | |
| | D | Mice no. | 8 | 13 | 15 | |
| | E | | 7 | 13 | 14 | |
| | F | | 7 | 13 | 15 | |
| | G | | 7 | 13 | 15 | |
| 3 | | Samples | $CR_{WT}$ | $CR_{M12}$ | | |
| | A | Mice no. | 13 | 15 | | |
| | B | | 13 | 14 | | |
| | C | | 11 | 14 | | |
| | D | | 3 | 3 | | |
| | E | | 3 | 3 | | |
| 4 | | Samples | UI | $CR_{WT}$ | $CR_{M12}$ | |
| | A | Mice no. | 8 | 13 | 15 | |
| | | Samples | $CR_{WT}$ | $CR_{M12}$ | | |
| | B | Mice no. | 11 | 12 | | |
| | C | | 11 | 12 | | |
| | | Samples | UI | $CR_{WT}$ | $CR_{M12}$ | |
| | D | Mice no. | 8 | 13 | 15 | |
| | E | | 8 | 13 | 15 | |
| 5 | | Samples | UI | $CR_{WT}$ | $CR_{M12}$ | |
| | A | Mice No. | 8 | 13 | 15 | |
| | B | | 7 | 13 | 14 | |
| | C | | 8 | 16 | 16 | |
| | D | | 8 | 13 | 15 | |
| S1 | | Samples | UI young | $CR_{WT}$ young | UI aged | $CR_{WT}$ aged |
| | A | Mice No. | 6 | 6 | 7 | 13 |
| | B | | 5 | 6 | 7 | 13 |
| | C | | 6 | 6 | 8 | 13 |
| | D | | 5 | 6 | 8 | 13 |
| S2 | | Samples | UI | $CR_{WT}$ | $CR_{M12}$ | |
| | A | | 7 | 13 | 14 | |
| S4 | | Samples | UI | $CR_{WT}$ | $CR_{M12}$ | |
| | A | Mice No. | 8 | 13 | 14 | Ifn-γ |
| | B | | 8 | 13 | 14 | Il-1β |
| | C | | 8 | 12 | 12 | Il-17A |
| | D | | 8 | 13 | 14 | Il-22 |
| | E | | 8 | 13 | 14 | Il-18 |
| | F | | 7 | 8 | 10 | Il-10 |
| | G | | 8 | 13 | 13 | Tnf |
| | H | | 8 | 11 | 13 | Il-6 |
| | I | | 8 | 13 | 13 | Cxcl1 |
| | J | | 8 | 12 | 14 | Ccl3 |
| | K | | 7 | 12 | 14 | Gm-csf |
| | L | | 8 | 13 | 14 | G-csf |

**Supplementary Table 2.**

Proteins differentially regulated between Epcam+/CR+ and Epcam+/CR- IECs

| Direction of change relative to UI | Epcam+/$CR_{WT}$+ | Epcam+/$CR_{WT}$- | No. of proteins |
|---|---|---|---|
| Opposite | Up | Down | 4 |
| | Up | No change | 75 |
| | Down | Up | 9 |
| | Down | No change | 86 |
| | No change | Up | 453 |
| | No change | Down | 101 |
| Same | Up | Up | 86 |
| | Down | Down | 28 |
| **Opposite direction differences** | | | **728** |
| **Same direction differences** | | | **114** |
| **Total differences** | | | **842** |

| Direction of change relative to UI | Epcam+/$CR_{M12}$+ | Epcam+/$CR_{M12}$- | No. of proteins |
|---|---|---|---|
| Opposite | Up | Down | 5 |
| | Up | No change | 9 |
| | Down | Up | 4 |
| | Down | No change | 35 |
| | No change | Up | 31 |
| | No change | Down | 38 |
| Same | Up | Up | 5 |
| | Down | Down | 3 |
| **Opposite direction differences** | | | **122** |
| **Same direction differences** | | | **8** |
| **Total differences** | | | **130** |

**Supplementary Table 3.**

Pathways specifically up- or downregulated in $Epcam^+/CR_{WT/M12}^+$ compared to $Epcam^+/CR_{WT/M12}^-$

| **Upregulated in $Epcam^+/CR_{WT}^+$ but downregulated or no changes in $Epcam^+/CR_{WT}^-$** | **Downregulated in $Epcam^+/CR_{WT}^+$ but upregulated or no changes in $Epcam^+/CR_{WT}^-$** |
|---|---|
| **Actin cytoskeleton/ pedestal and structural remodelling** | Actin cytoskeleton/ pedestal and structural remodelling |
| Coagulation/ ECM response | **Amino acid/ nitrogen metabolism** |
| **Epithelial surface/ mucosal surface** | Cell death and stress signalling |
| Innate immunity | Central carbon metabolism |
| **Ion transport** | Coagulation/ ECM response |
| **Ion/ redox/ miscellaneous metabolism** | DNA repair and genomic stability |
| Keratin/ epithelial stress program | Epithelial differeniation and identity |
| Lipid metabolism/ membrane remodelling | Glucose and nutrient transport |
| **Mitochondrial/ metabolic stress** | Immune sensors and innate defense |
| Proteostasis and stress response | **Ion transport** |
| Transcription/ signalling regulators | **Lipid and sterol transport** |
| Vesicle trafficking/endosomal system | Mitochondrial and oxidative metabolism |
| | NF-kB signalling pathway |
| | One-carbon and nucleotide metabolism |
| | Proliferation and cell cycle |
| | Proteostasis and stress response |
| | Proteostasis/Ubl conjugation pathway |
| | Redox and detox |
| | RNA processing |
| | **Stress-activated MAPK signalling** |
| | Transcriptional and epigenetic regulation |
| | **Tyrosine kinase signalling** |
| | Vesicle trafficking/endosomal system |

| **Upregulated in $Epcam^+/CR_{M12}^+$ but downregulated or no changes in $Epcam^+/CR_{M12}^-$** | **Downregulated in $Epcam^+/CR_{M12}^+$ but upregulated or no changes in $Epcam^+/CR_{M12}^-$** |
|---|---|
| Coagulation/acute phase response | Actin cytoskeleton/ pedestal and structural remodelling |
| Immunity | Apoptosis and cell survival signalling |
| Innate immunity | Carbohydrate metabolism |
| Keratin/ epithelial stress program | Coagulation/ECM response |
| Lipid metabolism | DNA repair and genomic stability |
| Proteostasis and stress response | Epithelial differeniation and identity |
| Transcriptional and epigenetic regulation | Glucose and nutrient transport |
| Vesicle trafficking/endosomal system | Innate immunity |
| | NF-kB signalling pathway |
| | One-carbon and nucleotide metabolism |
| | Proteostasis/Ubl-conjugation pathway |
| | Redox and detox |
| | RNA processing |
| | Transcriptional and epigenetic regulation |
| | **Translation machinery** |
| | Translation, RNA and transcription regulation |
| | Vesicle trafficking/endosomal system |

**Supplementary Table 4.**

Pathways specifically up or downregulated in $EpCAM^+/CR_{WT/M12}^-$ compared to $EpCAM^+/CR_{WT/M12}^+$

| **No changes in $Epcam^+/CR_{WT}^+$ but upregulated in $Epcam^+/CR_{WT}^-$** | **No changes in $Epcam^+/CR_{WT}^+$ but downregulated in $Epcam^+/CR_{WT}^-$** |
|---|---|
| **Amino acid and nutrient transport** | Actin dynamics, focal adhesion and contractility |
| **Apoptosis** | Cell adhesion and immune modulation |
| **Cell adhesion and cytoskeleton** | **Epithelial barrier and structural proteins** |
| Central carbon metabolism | Extracellular matrix, and stromal components |
| **Chromatin remodelling and epigenetic regulation** | Glycosylation and Mucin-type-O-Glycan synthesis |
| **DNA replication, repair and cell cycle** | Immunity |
| Epithelial barrier and structural proteins | **Innate immunity** |
| GPCR and second messanger signalling | Mitochondrial metabolism and bioenergetics |
| **Hormone and growth factor signalling** | Proteostasis and stress response |
| Innate immunity | **Redox and oxidative stress** |
| **Ion and solute transport** | **Signal transduction** |
| **Lipid metabolism** | Vesicle trafficking/endosomal system |
| Metabolism and mitochondria | Xenobiotic and lipid metabolism |
| mRNA processing, nuclear body organisation | |
| **NAD, PAR, Cell stress Signalling** | |
| Nuclear transport | |
| One-carbon and nucleotide metabolism | |
| Protein transport | |
| Proteostasis and stress response | |
| Redox and oxidative stress | |
| Ribosome biogenesis and Nucleolar Program | |
| RNA processing, splicing, and export | |
| **Secretory/enteroendocrine cells** | |
| **Stem/Regerative signalling** | |
| Transcription regulation | |
| Translation machinery | |
| Vesicle trafficking/endosomal system | |

| **No changes in $Epcam^+/CR_{M12}^+$ but upregulated in $Epcam/CR_{M12}^-$** | **No changes in $Epcam^+/CR_{M12}^+$ but downregulated in $Epcam^+/CR_{M12}^-$** |
|---|---|
| **cAMP/PKA signalling** | Actin dynamics, focal adhesion and contractility |
| Central carbon metabolism | Coagulation and complement system |
| Epithelial barrier integrity/keratinocyte differentiation | Extracellular matrix, and stromal components |
| GPCR and second messanger signalling | Glycosylation and Mucin-type-O-Glycan synthesis |
| Innate immunity | Immunity |
| Metabolism and mitochondria | Lipid transport and lipoprotein metabolism |
| mRNA processing, nuclear body organisation | Mitochondrial metabolism and bioenergetics |
| Nuclear transport | Proteostasis and stress response |
| One-carbon and nucleotide metabolism | Vesicle trafficking and membrane dynamics |
| Proteostasis and stress response | |
| Redox and oxidative stress | |
| Ribosome biogenesis and Nucleolar Program | |
| RNA processing, splicing, and export | |
| Transcriptional regulation | |
| Translation machinery | |
| Vesicle trafficking/endosomal system | |
